# The Two-Component System CrdRS Mediates Sub-inhibitory Antibiotic-Induced Biofilm Formation in Helicobacter pylori via Bidirectional Regulation of Transporters PlpA and GlnP

**DOI:** 10.64898/2026.06.29.735229

**Authors:** Wenxin Zhang, Lu Zhang, Yanlin Sun, Mingzhong Zhao, Shujing Xu, Han Yu, Wenjing Wang, Jinmeng Liu, Wenyue Ma, Xiaoyu Wang, Ziyan Zhang, Yundong Sun

**Affiliations:** Key Laboratory for Experimental Teratology of the Ministry of Education and Department of Microbiology, School of Basic Medical Sciences, Cheeloo College of Medicine, Shandong University, Jinan, Shandong, China; Department of Pathology, School of Basic Medical Sciences, Cheeloo College of Medicine, Shandong University, Jinan, Shandong, China

**Keywords:** *Helicobacter pylori*, biofilm, two-component system CrdRS, sub-inhibitory antibiotics, ABC transporter, reactive oxygen species

## Abstract

Biofilm formation by *Helicobacter pylori* is a major driver of antibiotic tolerance and treatment failure, yet the signaling pathways that trigger biofilm development under sub-inhibitory antibiotic pressure remain poorly understood. Here, we show that sub-MIC levels of metronidazole, amoxicillin, and ciprofloxacin potently induce dense *H. pylori* biofilms. Through transcriptomic and genetic analyses, we identify the two-component system CrdRS as a central signaling hub that orchestrates this response via the bidirectional regulation of two ATP-binding cassette (ABC) transporters. Phosphorylated CrdR binds to AC-rich promoter motifs to directly activate *plpA*, which encodes a substrate-binding protein that drives exopolysaccharide secretion and matrix assembly. Concomitantly, CrdR represses *glnP*, which encodes an inner-membrane permease, thereby relieving transcriptional inhibition of the L-asparaginase gene *ansB*. This derepression triggers aberrant reactive oxygen species (ROS) accumulation, which promotes oxidative stress-dependent biofilm maturation. Through phenotypic analysis of c*rdRS* deletion mutants, phosphorylation-defective point mutants (CrdR^D53A^ and CrdS^H173A^), and exogenous hydrogen peroxide (H₂O₂) induction, we demonstrate that CrdRS specifically responds to antibiotic-induced stress signals in a manner genetically separable from ROS sensing. Collectively, our findings establish a dual-mechanism model in which CrdRS orchestrates antibiotic-induced biofilm formation by simultaneously controlling matrix production and intracellular ROS generation. Notably, *glnP* expression was significantly lower in clinical multidrug-resistant isolates than in drug-sensitive ones, whereas *plpA* showed the opposite trend. These findings provide a mechanistic foundation for developing CrdRS-targeted strategies to combat biofilm-associated *H. pylori* infections.

## Introduction

*Helicobacter pylori*, a gastric pathogen that persistently colonizes approximately half of the global population, is etiologically linked to a spectrum of gastroduodenal diseases, including chronic gastritis, peptic ulcer disease, mucosa-associated lymphoid tissue lymphoma, and gastric adenocarcinoma (1). Current clinical management relies on combination antibiotic regimens, typically comprising a proton pump inhibitor plus two or three antimicrobial agents (2). However, the relentless rise of antibiotic resistance has steadily eroded eradication success rates (3). Global surveillance data indicate that primary resistance to clarithromycin, metronidazole, and levofloxacin has reached alarming levels in many regions; clarithromycin resistance alone exceeds 15% in most WHO regions, rendering standard triple therapy unacceptably ineffective (4, 5).

Beyond genetically encoded resistance, a growing body of clinical evidence implicates biofilm formation as a critical factor in treatment failure. In patients who have failed multiple eradication regimens, *H. pylori* is frequently found organized into dense biofilms on the gastric mucosa, and isolates from such patients exhibit significantly higher biofilm-forming capacity than those from treatment-naïve individuals (6, 7). Biofilms are surface-associated bacterial communities encased in a self-produced matrix of extracellular polymeric substances (EPS). This matrix acts as a physical diffusion barrier and creates metabolically dormant microenvironments, conferring phenotypic tolerance that can exceed planktonic susceptibility by 10- to 1000-fold (8). Biofilm-mediated tolerance thus constitutes a major obstacle to successful eradication and a key driver of persistent infection (9, 10) .

*H. pylori* biofilm development proceeds through sequential stages: initial attachment, microcolony formation, maturation, and dispersal (11). Planktonic cells first adhere to gastric epithelium or abiotic surfaces through flagellar motility and outer membrane adhesins such as BabA, SabA, and HopQ (12–14); deletion of motility genes—*motB*, *fliM*, and *fliA*—severely impairs biofilm initiation. Mature biofilms are structurally heterogeneous, comprising spiral and coccoid forms embedded in an EPS matrix rich in proteomannans, fucose, galactose, *N*-acetylglucosamine, eDNA, and proteins. The quorum-sensing molecule AI-2, synthesized by LuxS, modulates biofilm dynamics in a complex, concentration-dependent manner: paradoxically, luxS deletion mutants incapable of AI-2 production exhibit enhanced biofilm formation, suggesting that AI-2 signaling helps regulate the planktonic-to-biofilm transition (15). Despite substantial progress in characterizing *H. pylori* biofilm structure and composition, the upstream signals and signaling pathways that trigger this transition remain largely undefined (8).

In the gastric niche, *H. pylori* faces continuous environmental stress—reactive oxygen and nitrogen species generated by host inflammatory responses, pH fluctuations, and nutrient limitation (11, 16, 17). Under such adverse conditions, adopting a biofilm lifestyle represents a critical adaptive strategy that enhances bacterial survival (8). During antibiotic therapy, pharmacokinetic constraints—including limited drug penetration through the gastric mucus layer, acid-catalyzed degradation, and uneven tissue distribution—often prevent sustained bactericidal concentrations at the infection site, creating exposure windows at sub-MICs (18). Accumulating evidence indicates that sub-MIC antibiotics act not merely as ineffective antimicrobials, but as potent environmental signals that trigger bacterial stress responses and induce biofilm formation (19, 20). In *H. pylori*, Krzyżek et al. recently showed that continuous exposure to sub-MICs of metronidazole or levofloxacin accelerates autoaggregation, enhances EPS production, and increases biofilm dimensions, whereas clarithromycin exerts the opposite effect (21). This phenomenon mirrors observations in other clinically significant pathogens—*Staphylococcus aureus*, *Pseudomonas aeruginosa*, and *Escherichia coli*—where sub-lethal antibiotic concentrations promote biofilm development, suggesting a phylogenetically conserved adaptive strategy (22, 23). Nevertheless, the molecular machinery—specifically, the sensory systems and downstream regulatory cascades—that enables *H. pylori* to perceive sub-MIC antibiotics as signals and translate them into a biofilm developmental program has yet to be systematically elucidated.

Bacteria rely on two-component systems (TCS) as primary signal-transduction mechanisms for sensing and responding to environmental changes. A canonical TCS consists of a membrane-anchored sensor histidine kinase (HK) and a cognate cytoplasmic response regulator (RR). Upon stimulus perception, the HK undergoes ATP-dependent autophosphorylation at a conserved histidine residue and subsequently transfers the phosphoryl group to an aspartate residue on the RR. Phosphorylation induces a conformational change that activates the RR’s DNA-binding domain, enabling sequence-specific promoter recognition and target gene modulation. The *H. pylori* genome encodes a limited TCS repertoire that governs critical adaptive responses: ArsRS senses acidic pH and regulates acid-adaptive genes, including the urease cluster (24); FlgRS controls flagellar biosynthesis and motility (25); and CrdRS (HP1364/HP1365) was originally characterized as a copper-sensing system that maintains metal ion homeostasis by activating the copper resistance gene *crdA* (26) and was later shown to also respond to nitrosative stress (27). Notably, several TCSs have been implicated in biofilm regulation. ArsRS negatively regulates biofilm development: *arsS* deletion upregulates the outer membrane protein HomB and enhances biofilm formation (28). FlgR, via inhibition of the alternative sigma factor σ⁵⁴, represses the molybdate transport system ModABD, thereby suppressing biofilm formation; ROS-induced allosteric inhibition of the nickel-responsive regulator NikR relieves this FlgR-mediated repression, providing a mechanistic link between oxidative stress and biofilm induction (29). These findings establish that TCSs can integrate diverse environmental cues—pH, metal ions, oxidative stress—to control the biofilm developmental switch. However, whether any TCS senses antibiotic signals to mediate sub-MIC-induced biofilm formation has remained an open question.

In this study, we systematically investigated the signaling network underlying sub-MIC antibiotic-induced biofilm formation in *H. pylori*. We demonstrate that CrdRS functions as a central antibiotic-sensing hub that orchestrates biofilm development through a dual regulatory mechanism: phosphorylated CrdR directly activates *plpA* to drive exopolysaccharide secretion and matrix assembly, while concurrently repressing *glnP*, thereby triggering *ansB*-dependent ROS accumulation and oxidative stress-driven biofilm maturation.

## Results

### Sub-inhibitory concentrations of antibiotics induce biofilm formation in *H. pylori* in vitro

To determine whether low antibiotic concentrations induce biofilm formation in *H. pylori*, we selected three clinically relevant antibiotics—metronidazole (MTZ), amoxicillin (AMO), and ciprofloxacin (CIP). For each, we established three concentration gradients—two sub-MIC levels and one at the MIC (Figure 1)—and examined their effects on biofilm formation by the wild-type strain 26695 (WT) using confocal laser scanning microscopy (CLSM) and scanning electron microscopy (SEM). Sub-inhibitory metronidazole at 1 μg/mL induced the most robust biofilm formation, comparable to that elicited by low-concentration H₂O₂ (positive control). CLSM revealed that biofilms formed under 1 μg/mL metronidazole exhibited a highly compact architecture with markedly reduced microchannels and interbacterial spaces, and were substantially thicker than those formed under other conditions (Figure 1a). SEM confirmed that the extracellular matrix of these biofilms was densely organized (Figure 1b). As metronidazole concentration increased, biofilm biomass decreased significantly; at the MIC, biofilm thickness was markedly reduced (Figure 1a) and matrix content substantially diminished (Figure 1b). Similarly, sub-MICs of AMO (1/32 μg/mL) and CIP (1/16 μg/mL) both induced dense, intact biofilms (Figures S1 and S2).

**Figure 1.**
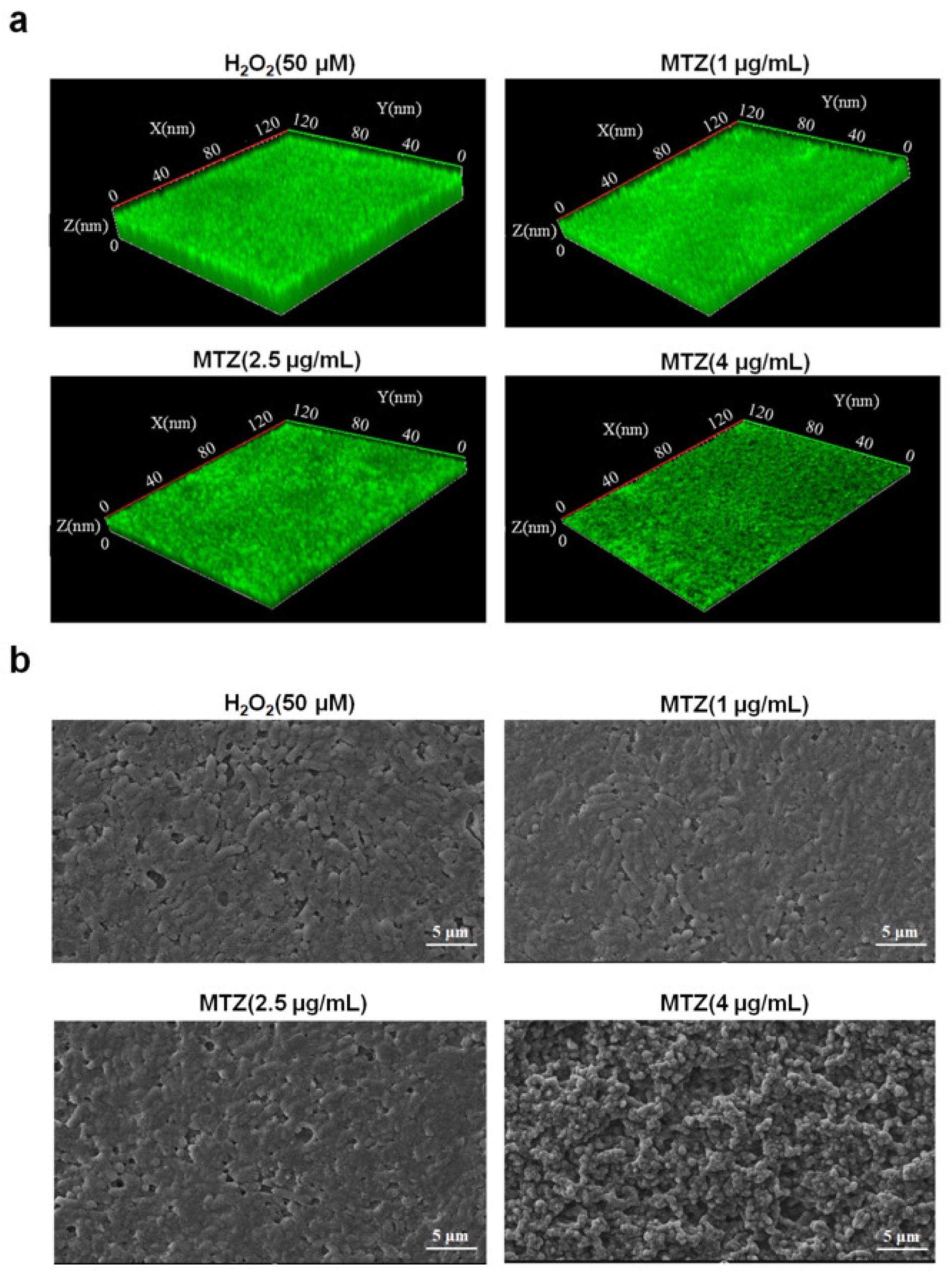
Sub-inhibitory concentrations of metronidazole induce *H. pylori* biofilm formation. Wild-type (WT) *H. pylori* strain 26695 was cultured on nitrocellulose membranes for 72 h in the presence of metronidazole at the indicated concentrations (0, 1, 2.5, and 4 μg/mL). Untreated WT (0 μg/mL) served as the negative control, and WT exposed to 50 μM H₂O₂ served as the positive control. (a) Representative confocal laser scanning microscopy (CLSM) images of biofilms stained with SYTO™ 9 (green) to visualize viable bacteria. (b) Representative scanning electron microscopy (SEM) images showing biofilm ultrastructure. Scale bar, 5 μm. Data are representative of three independent experiments.

### PlpA is essential for sub-inhibitory antibiotic-induced biofilm formation

To elucidate the molecular mechanisms underlying sub-inhibitory antibiotic-induced biofilm formation, we performed transcriptomic analysis on WT biofilm cells treated with the three antibiotics. KEGG pathway enrichment revealed that differentially expressed genes were significantly enriched in “ABC transporters” and “alanine, aspartate, and glutamate metabolism.” Notably, the outer-membrane substrate-binding protein gene *plpA* (*hp1564*) was markedly upregulated, whereas the inner-membrane amino acid permease gene cluster *glnP* (*hp1169*–*hp1170*) was significantly downregulated (Figure S3a). Clustering heatmap analysis further confirmed consistent elevation of *plpA* expression across all treatment groups, with *glnP* showing the opposite trend (Figure S3b).

To verify whether PlpA mediates sub-inhibitory antibiotic-induced biofilm formation, we constructed a *plpA* deletion mutant (*ΔplpA*) and its complementation strain (*plpA*\*). Growth curve analysis revealed no significant growth defect in the *ΔplpA* strain (Figure S4a). After 72 h of induction with sub-inhibitory antibiotics, we assessed biofilm formation by CLSM and SEM. The WT and *plpA*\* strains formed structurally compact, intact biofilms with abundant extracellular matrix, whereas the *ΔplpA* strain displayed virtually no intact biofilm architecture, reduced extracellular matrix, and increased inter-membrane void spaces (Figure 2). Similar results were obtained with amoxicillin and ciprofloxacin (Figures S5 and S6). Moreover, the *plpA* deletion mutant in the clinical isolate H57 (H57*ΔplpA*) exhibited markedly attenuated biofilm-forming capacity compared with the H57 WT strain (Figure S7).

**Figure 2.**
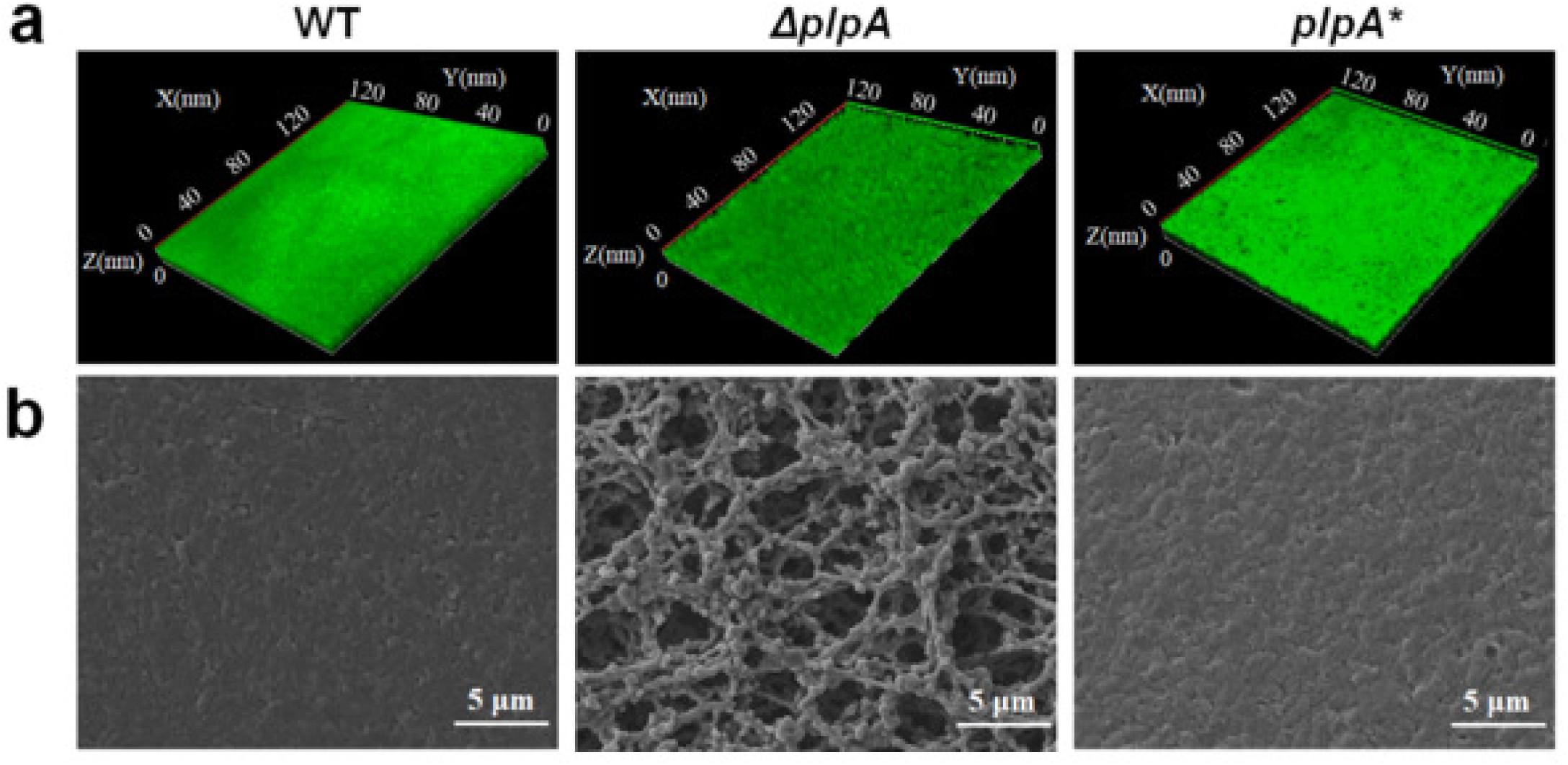
PlpA is essential for sub-inhibitory metronidazole-induced biofilm formation in *H. pylori*. WT, *ΔplpA* and *plpA\** complemented strains were cultured on nitrocellulose membranes containing metronidazole (1 μg/mL) for 3 days to allow mature biofilm development. (a) Representative CLSM images of biofilms stained with SYTO™ 9 (green). (b) Representative SEM images of biofilm architecture. Scale bar, 5 μm. Data are representative of three independent experiments.

### Downregulation of GlnP mediates biofilm formation under sub-inhibitory antibiotic stress

To investigate whether GlnP participates in biofilm regulation, we generated a *glnP* deletion mutant (*ΔglnP*) and its complementation strain (*glnP\**). Growth curve analysis showed that the mutant and complementation strains grew comparably to WT (Figure S4b). Following 24 h of treatment with sub-inhibitory metronidazole (1 μg/mL), we assessed biofilm morphology. Compared with WT and *glnP\**, the *ΔglnP* mutant formed substantially denser and thicker biofilms (Figure 3a) with increased extracellular matrix content (Figure 3b). Similar phenotypes were observed with amoxicillin and ciprofloxacin (Figures S8 and S9). Furthermore, deletion of *glnP* in the clinical isolate H57 (H57*ΔglnP*) led to markedly enhanced biofilm formation compared with the parental H57 strain (Figure S10).

**Figure 3.**
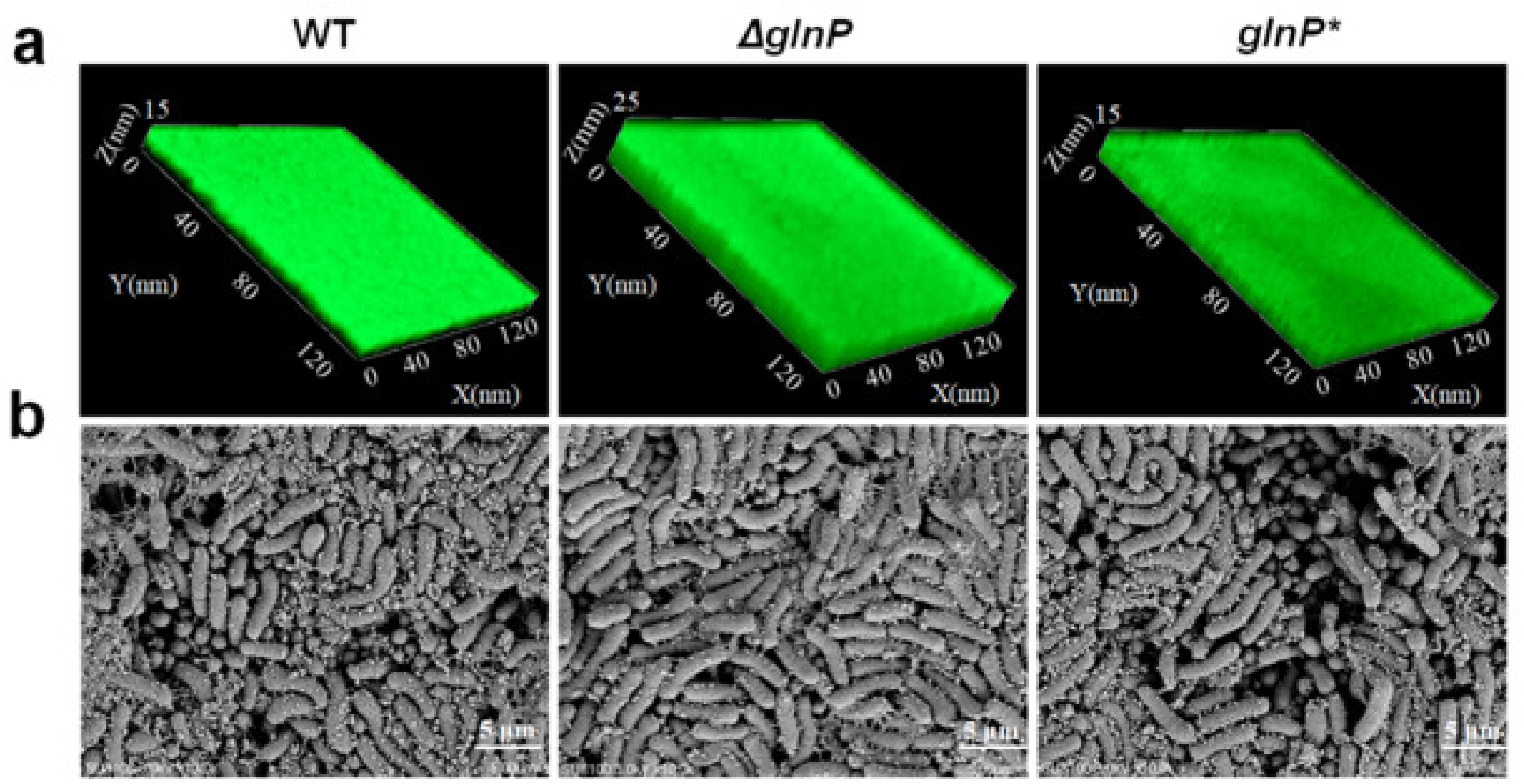
Deletion of *glnP* enhances sub-inhibitory metronidazole-induced biofilm formation in *H. pylori*.WT, *ΔglnP* and *glnP\** complemented strains were cultured on nitrocellulose membranes containing metronidazole (1 μg/mL) for 3 days. (a) Representative CLSM images of biofilms stained with SYTO™ 9 (green). (b) Representative SEM images showing biofilm ultrastructure. Scale bar, 5 μm. Data are representative of three independent experiments.

### The CrdRS two-component system is required for sub-inhibitory antibiotic-induced biofilm formation

To explore how *plpA* expression is elevated and *glnP* expression suppressed during biofilm formation, we performed transcriptomic analysis to identify differentially expressed regulatory genes in biofilm cells. Hierarchical clustering and heatmap visualization revealed marked upregulation of the transcriptional regulator gene *hp1365* (Figure S11). *hp1365* encodes CrdR, the response regulator of the CrdRS two-component system (26). Given this high expression and the established signaling roles of TCSs, we hypothesized that CrdRS participates in antibiotic-induced biofilm formation.

To test this, we constructed crdRS deletion mutants (*ΔcrdR* and *ΔcrdS*), complementation strains (*crdR\** and *crdS\**), and phosphorylation-defective point mutants (CrdR^D53A^and CrdS^H173A^). All strains exhibited growth kinetics comparable to WT (Figures S4c–e). Following induction with sub-inhibitory antibiotics, we examined biofilm morphology. The *ΔcrdR*, *ΔcrdS*, CrdR^D53A^, and CrdS^H173A^ strains all showed reduced biofilm thickness and biomass, increased intercellular microchannels and void spaces, rougher surface textures, and diminished extracellular matrix (Figures 4, 5, and S12–S15). In contrast, the complementation strains formed intact biofilms comparable to those of WT (Figures 4, 5).

**Figure 4.**
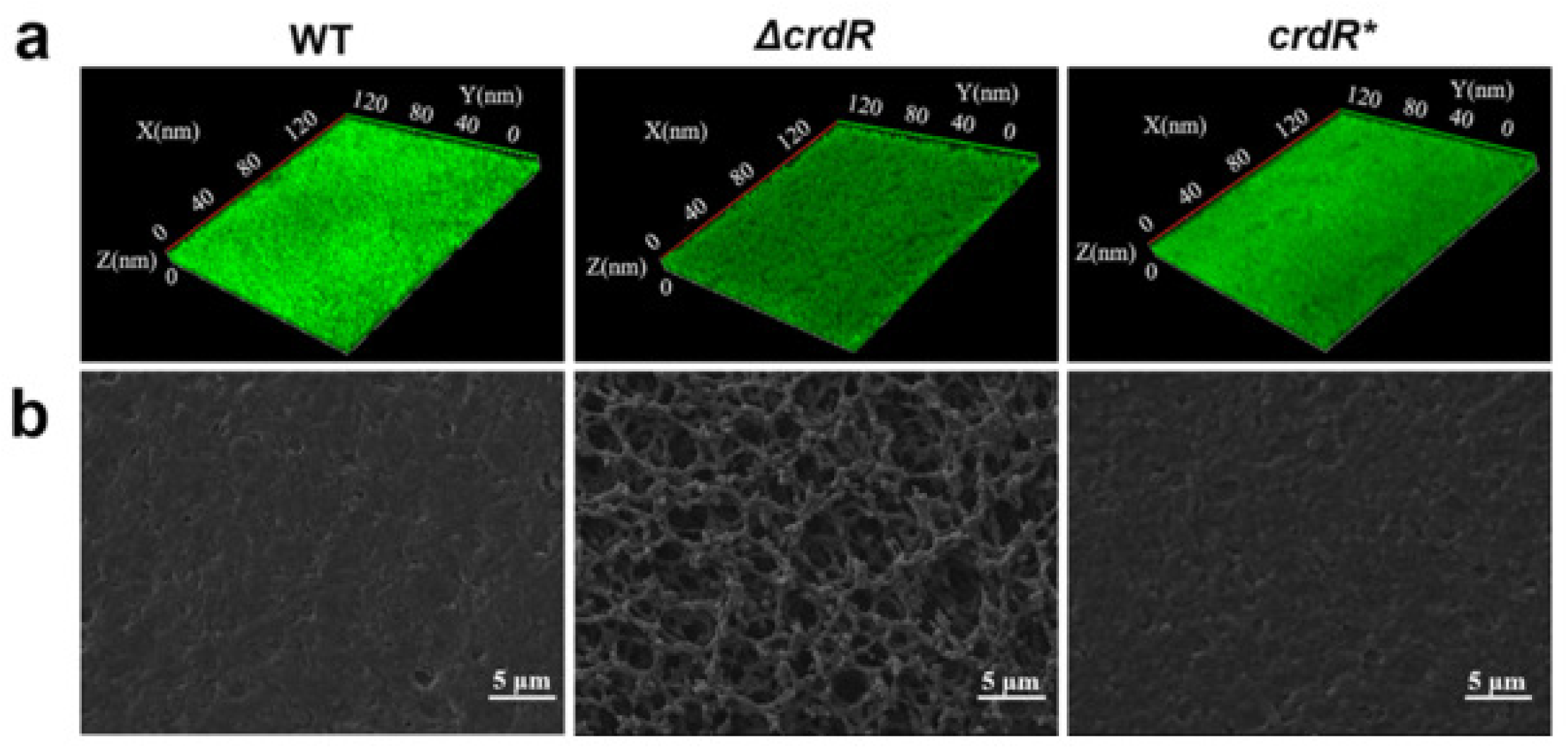
CrdR is required for sub-inhibitory metronidazole-induced biofilm formation in *H. pylori*. WT, *ΔcrdR* and *crdR\**complemented strains were cultured on nitrocellulose membranes containing metronidazole (1 μg/mL) for 3 days. (a) Representative CLSM images of biofilms stained with SYTO™ 9 (green). (b) Representative SEM images of biofilm architecture. Scale bar, 5 μm. Data are representative of three independent experiments.

**Figure 5.**
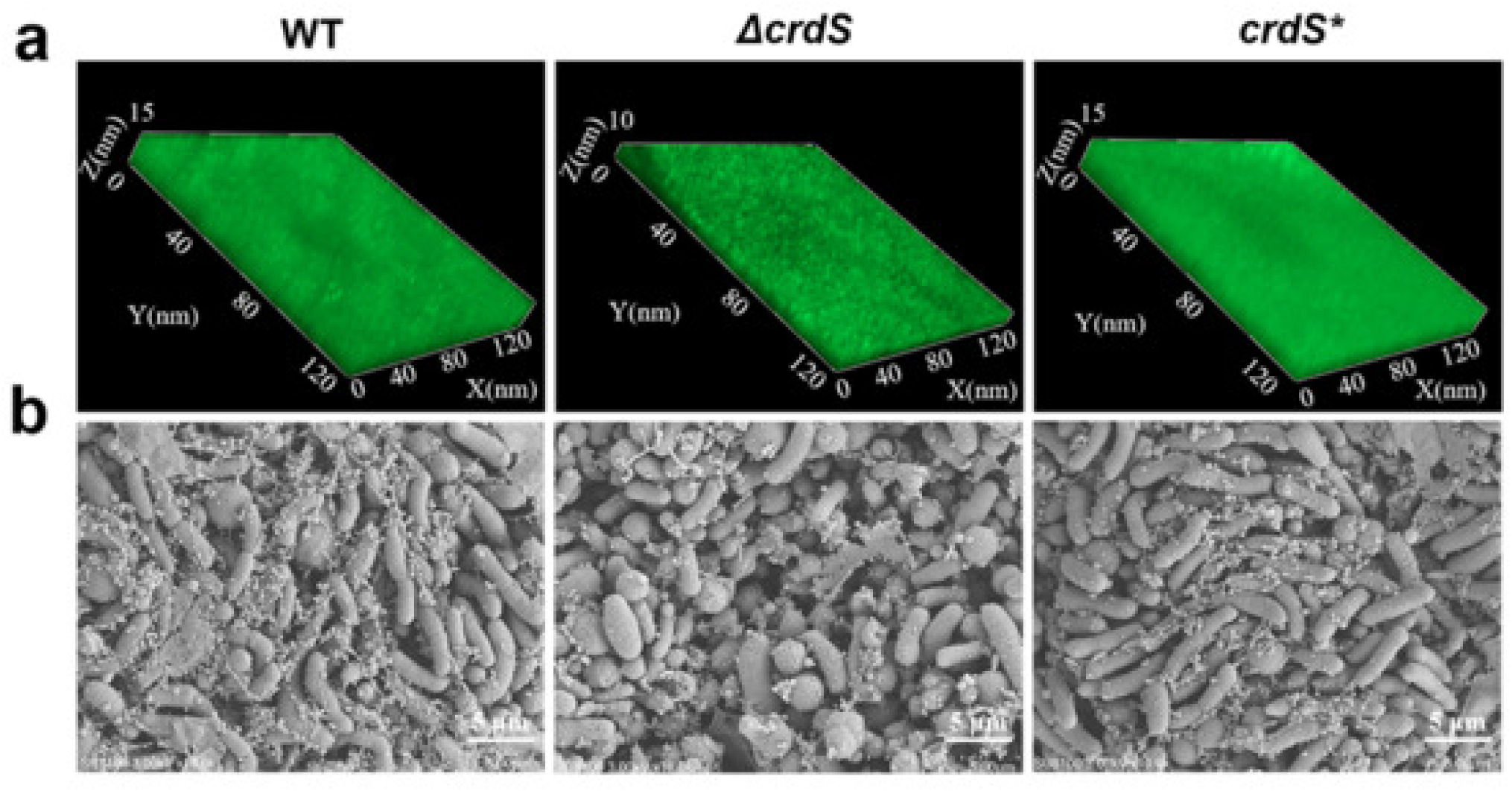
CrdS is required for sub-inhibitory metronidazole-induced biofilm formation in *H. pylori*. WT, *ΔcrdS* and *crdS\** complemented strains were cultured as described in Figure 4. (a) Representative CLSM images of biofilms stained with SYTO™ 9 (green). (b) Representative SEM images of biofilm ultrastructure. Scale bar, 5 μm. Data are representative of three independent experiments.

### CrdR upregulates the ABC transporter PlpA to promote antibiotic-induced biofilm formation

To verify whether CrdR regulates PlpA, we used qRT-PCR and semi-quantitative RT-PCR to measure *plpA* mRNA levels in biofilm cells of the *ΔcrdR* and *crdR\** strains. *plpA* expression was markedly reduced in the *ΔcrdR* strain relative to WT (Figures 6a, b), indicating that CrdR positively regulates *plpA* transcription. A luciferase reporter assay further confirmed this regulation. *E. coli* co-transformed with pGL3-OP*_plpA_* and pET32a-*crdR* exhibited significantly higher luciferase activity than cells harboring pGL3-OP*_plpA_* alone (Figures 6c, d), showing that CrdR activates *plpA* expression.

**Figure 6.**
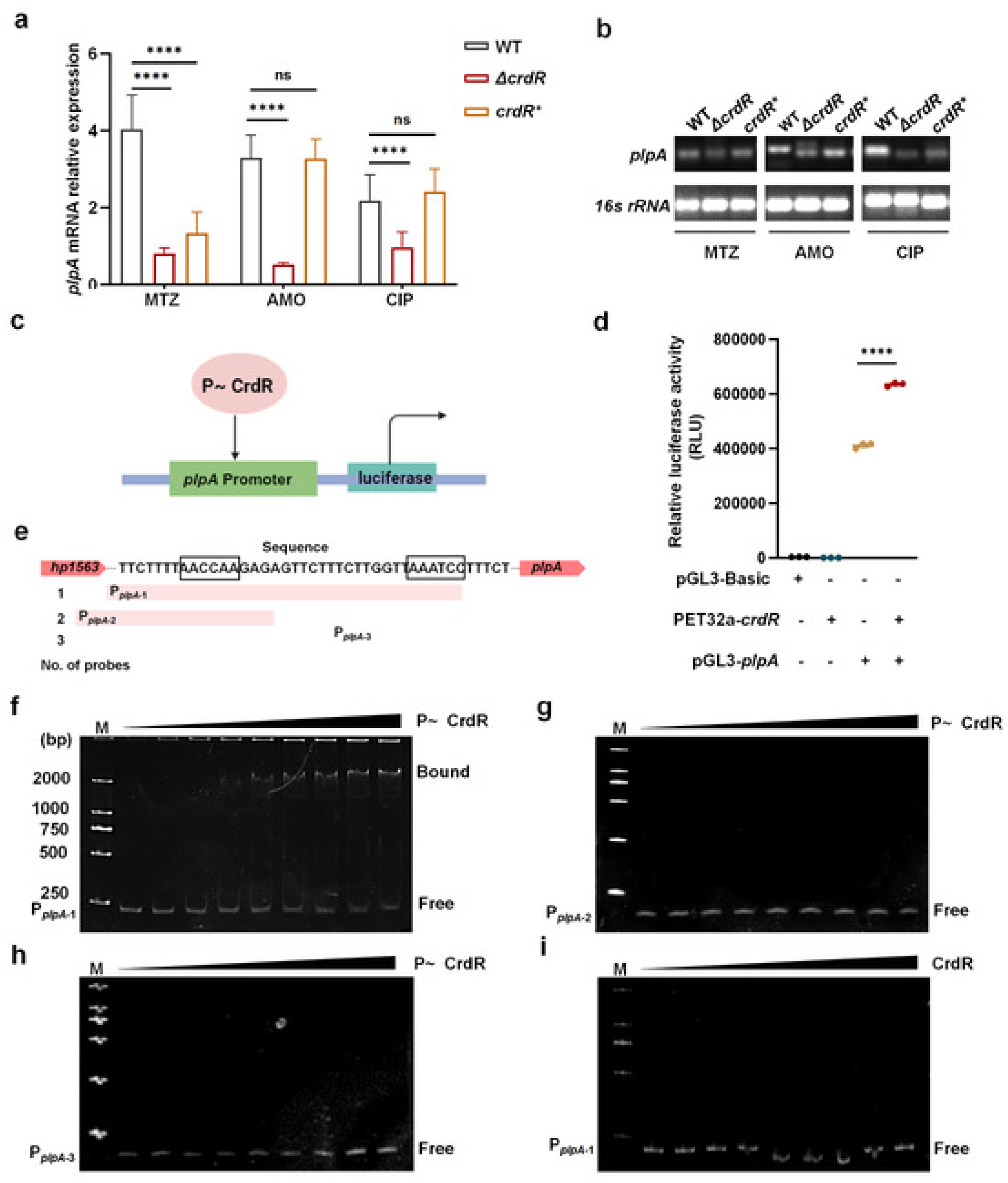
Phosphorylated CrdR directly binds the *plpA* promoter to activate its transcription. (a) Relative *plpA* mRNA levels in WT, *ΔcrdR*, and *crdR\** strains following treatment with sub-inhibitory concentrations of metronidazole (MTZ; 1 μg/mL), amoxicillin (AMO; 1/32 μg/mL), or ciprofloxacin (CIP; 1/16 μg/mL), determined by qRT-PCR. (b) Semi-quantitative RT-PCR analysis of *plpA* expression in the indicated strains. (c) Schematic of the luciferase reporter assay. (d) Luciferase activity in *E. coli* BL21 co-transformed with the reporter plasmid pGL3-OP*_plpA_* and the CrdR expression plasmid pET32a-*crdR* in the combinations indicated. Empty pGL3-Basic and pET32a vectors served as controls. (e) Schematic representation of EMSA probes spanning the *plpA* promoter region. The intergenic region between *hp1563* and *plpA* contains two AC-rich motifs (shaded boxes). Probe P*_plpA_*_-1_ (∼100 bp) encompasses both motifs; probes P*_plpA_*_-2_ and P*_plpA_*_-3_ (∼70 bp each) contain only the upstream or downstream motif, respectively. (f-i) Electrophoretic mobility shift assay (EMSA) analysis of CrdR binding. (f) Phosphorylated CrdR (P∼CrdR) binds to probe P*_plpA_*_-1_ containing dual AC-rich motifs, producing a shifted complex (Bound). (g) Unphosphorylated CrdR does not bind to this probe. (h, i) Phosphorylated CrdR does not bind to probes P*_plpA_*_-2_ or P*_plpA_*_-3_ containing single AC-rich motifs. Free probe, unbound DNA. Protein concentrations: 0, 1.5, 3, 4.5, 6, 7.5, 9, 10.5, and 12 μM. Data in (a) and (d) are presented as mean ± SD; ****, *P* < 0.0001; ns, not significant. All experiments were performed with three independent biological replicates.

We next investigated whether phosphorylated CrdR directly binds the *plpA* promoter using EMSA. We designed three DNA probes: probe P*_plpA_*_-1_ containing two AC-rich regions (∼100 bp); probe P*_plpA_*_-2_ containing the upstream AC-rich region (∼70 bp); and probe P*_plpA_*_-3_ containing the downstream AC-rich region (∼70 bp) (Figure 6e). Phosphorylated CrdR bound to probe P*_plpA_*_-1_, which harbors dual AC-rich motifs (Figure 6f), but did not bind to probes P*_plpA_*_-2_ or P*_plpA_*_-3_, which contain a single AC-rich region (Figures 6g, h). Unphosphorylated CrdR failed to bind any probe (Figure 6i). These results demonstrate that phosphorylated CrdR specifically binds the AC-rich region of the *plpA* promoter to activate transcription.

### CrdR downregulates the ABC transporter GlnP to promote antibiotic-induced biofilm formation

To assess whether CrdR regulates GlnP, we used qRT-PCR and RT-PCR to measure *glnP* (*hp1169*–*hp1170*) mRNA expression in *ΔcrdR* biofilm cells. *glnP* expression was markedly upregulated in the *ΔcrdR* strain compared with WT (Figures 7a, b), indicating that CrdR negatively regulates *glnP*. Luciferase reporter analysis further showed that *E. coli* co-transformed with pGL3-OP*_glnP_* and pET32a-*crdR* exhibited significantly reduced luciferase activity relative to cells expressing pGL3- OP*_glnP_* alone (Figures 7c, d), confirming that CrdR represses *glnP* expression.

**Figure 7.**
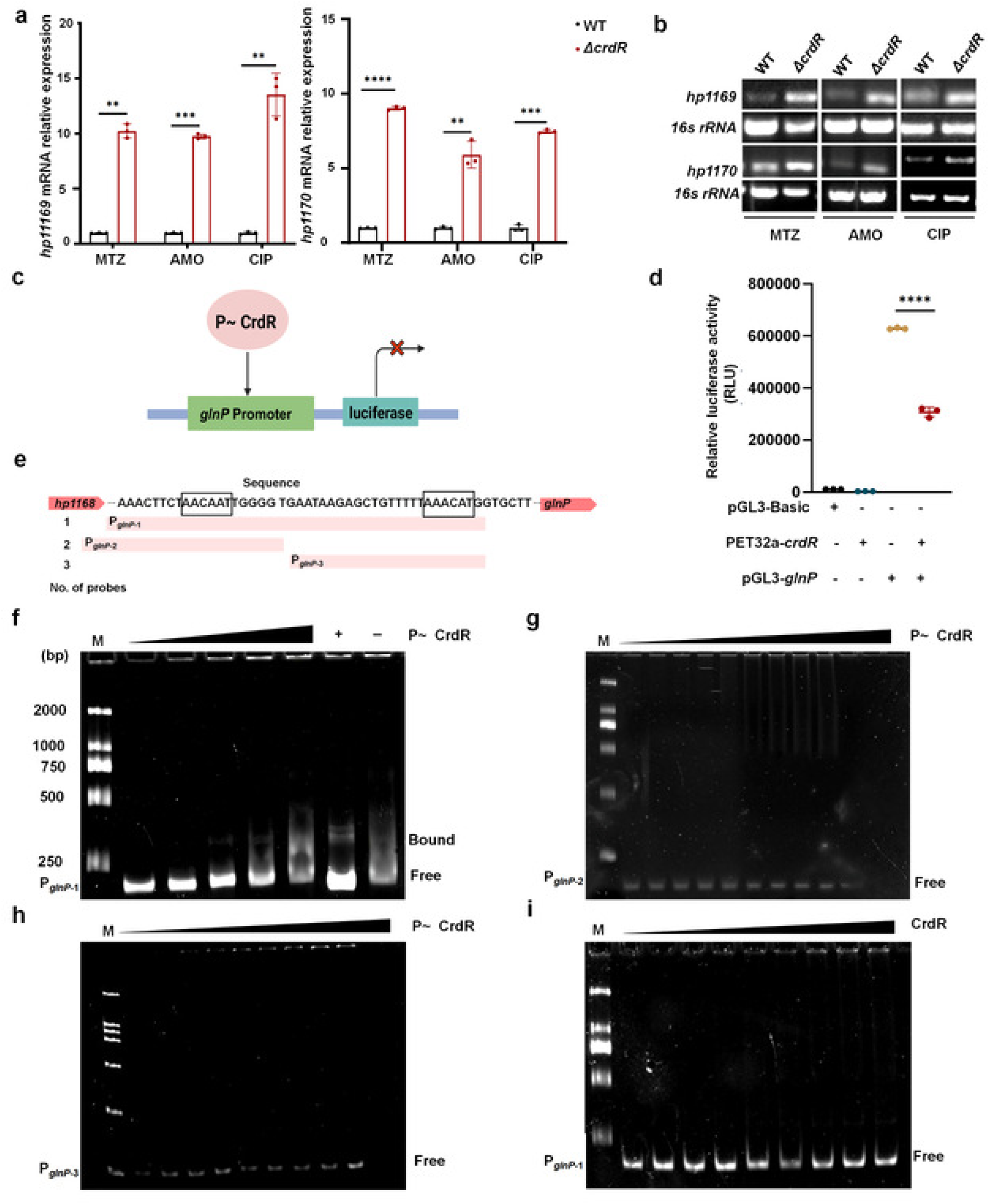
Phosphorylated CrdR directly binds the *glnP* promoter to repress its transcription. (a) Relative expression of the *glnP* gene cluster (*hp1169*-*hp1170*) in WT and *ΔcrdR* strains treated with sub-inhibitory MTZ, AMO, or CIP, determined by qRT-PCR. (b) Semi-quantitative RT-PCR analysis of *glnP* expression in the *ΔcrdR* strain. (c) Schematic of the luciferase reporter assay. (d) Luciferase activity in *E. coli* BL21 co-transformed with pGL3-OP*_glnP_* and pET32a-*crdR* in the indicated combinations. (e) Schematic representation of EMSA probes spanning the *glnP* promoter. Blue boxes indicate putative CrdR-binding sites. Probe P*_glnP_*_-1_ contains dual binding sites; probes P*_glnP_*_-2_ and P*_glnP_*_-3_ contain only the upstream or downstream single site, respectively. (f-i) EMSA analysis of CrdR binding to the *glnP* promoter. (f) Phosphorylated CrdR (P∼CrdR) specifically binds probe P*_glnP_*_-1_ containing dual sites. lane 6, *crdA* promoter probe (positive control); lane 7, random sequence probe (negative control). Protein concentrations: 0, 3, 6, 9, 11.5, 11.5, and 11.5 μM (g) Unphosphorylated CrdR does not bind. (h,i) P∼CrdR does not bind to probes P*_glnP_*_-2_ or P*_glnP_*_-3_ containing single sites. Bound, DNA-protein complex; Free probe, unbound DNA. (g,h,i)Protein concentrations: 0, 1.5, 3, 4.5, 6, 7.5, 9, 10.5, and 12 μM. Data in (a) and (d) are presented as mean ± SD; **, *P* < 0.01; ***, *P* < 0.001;****, *P* < 0.0001; ns, not significant. All experiments were performed with three independent biological replicates.

We next performed EMSA to test direct binding. We designed three probes: P*_glnP_*_-1_ containing two AC-rich regions (∼180 bp); P*_glnP_*_-2_ containing the upstream AC-rich region (∼120 bp); and P*_glnP_*_-3_ containing the downstream AC-rich region (∼70 bp) (Figure 7e). Phosphorylated CrdR specifically bound probe P*_glnP_*_-1_ containing dual AC-rich motifs (Figure 7f), but showed no significant interaction with the single-site probes (Figures 7g, h). Unphosphorylated CrdR did not bind (Figure 7i). Furthermore, the phosphorylation-site mutant CrdR^D53A^ was unable to form complexes with either the *glnP* or *plpA* dual-site probes, confirming that phosphorylation at Asp53 is essential for DNA-binding activity (Figure S16).

### Deletion of *plpA* reduces polysaccharide content in the biofilm extracellular matrix

To determine why *plpA* deletion impairs extracellular matrix formation, we compared the major EPS components—proteins, eDNA/RNA, and polysaccharides—in WT, *ΔplpA*, and *plpA*\* biofilms by specific staining. Polysaccharide content was markedly reduced in the *ΔplpA* biofilm EPS compared with WT, whereas eDNA/RNA content was only slightly decreased. The EPS composition of the *plpA*\*strain was comparable to that of WT (Figure 8).

**Figure 8.**
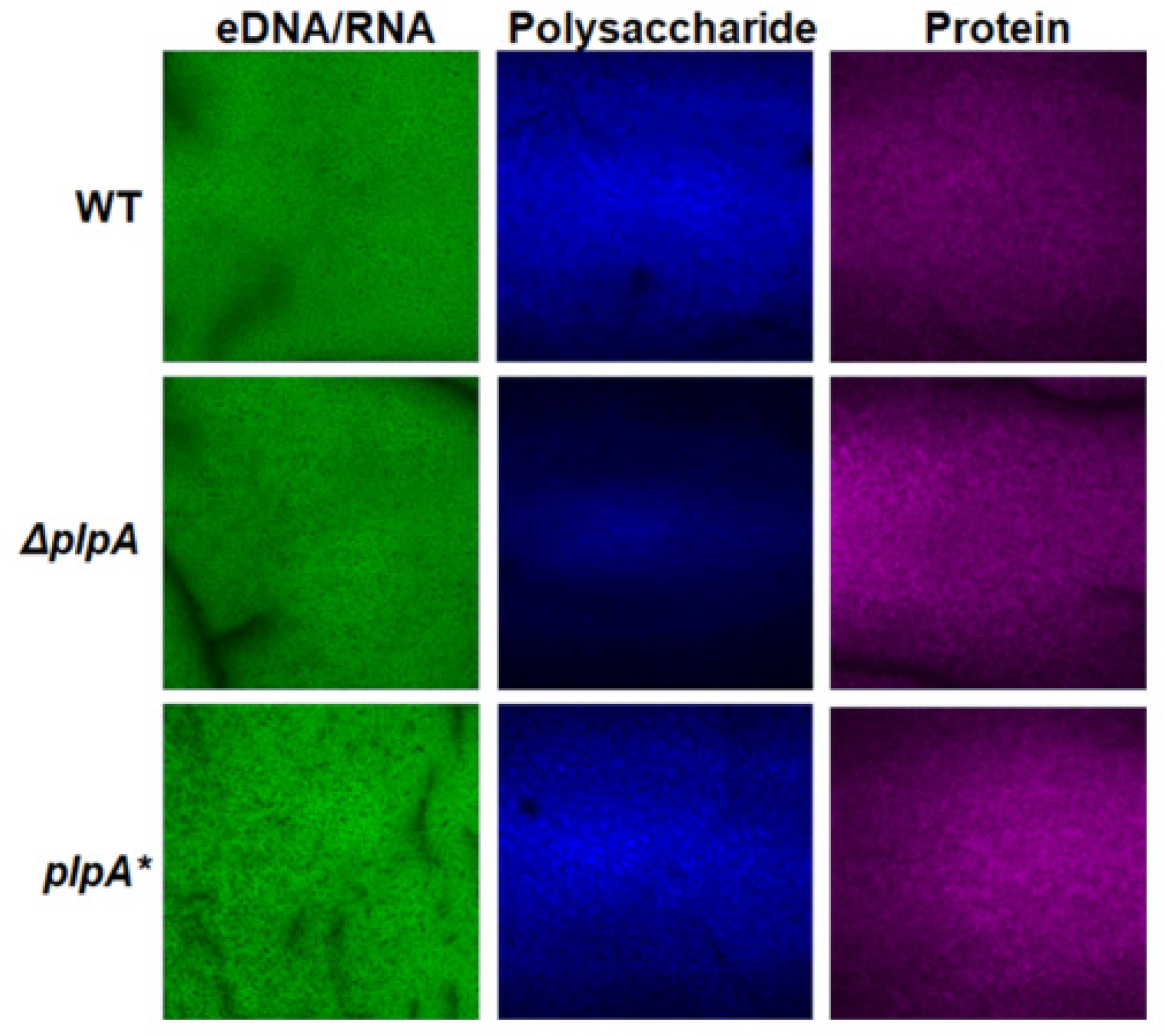
Deletion of *plpA* reduces polysaccharide content in the biofilm extracellular matrix. WT, *ΔplpA*, and *plpA\** biofilms were cultured on nitrocellulose membranes for 3 days and stained for extracellular polymeric substance (EPS) components. eDNA/RNA was visualized with propidium iodide (PI), exopolysaccharides with Calcofluor White, and extracellular proteins with SYPRO Ruby. Representative CLSM images are shown. Data are representative of three independent experiments.

### *glnP* deletion upregulates *ansB*, driving ROS-dependent biofilm formation

To elucidate the metabolic mechanisms through which *glnP* deletion promotes biofilm formation, we performed transcriptomic (RNA-seq) analysis on WT and *ΔglnP* strains under sub-inhibitory antibiotic stress. KEGG enrichment analysis showed that upregulated genes in the *ΔglnP*strain were significantly enriched in TCA cycle and amino acid metabolism pathways (Figure S17a). Notably, the L-asparaginase gene *ansB* (*hp0723*), which links these two metabolic pathways, was markedly upregulated (Figure S17b). Aspartate generated by AnsB-catalyzed deamination can enter the TCA cycle via transamination, consuming α-ketoglutarate (α-KG), a core precursor for glutathione (GSH) synthesis. Depletion of the α-KG pool impairs antioxidant capacity, leading to intracellular ROS accumulation(30). Given that low-level ROS are known to induce biofilm formation in *H. pylori* (16), we hypothesized that *glnP* deletion activates *ansB*, causing oxidative stress and ROS-dependent biofilm assembly.

To verify the role of *ansB*, we constructed a *ΔglnPΔansB* double mutant. Growth curve analysis showed that the double mutant grew comparably to WT and the *ΔglnP* single mutant (Figure S4f). We compared biofilm formation induced by sub-inhibitory concentrations of metronidazole, amoxicillin, and ciprofloxacin among WT, *ΔglnP,* and *ΔglnPΔansB* strains. The *ΔglnPΔansB* strain exhibited significantly reduced biofilm thickness, loosely arranged bacterial architecture, and diminished EPS compared with the *ΔglnP* strain, indicating that deletion of *ansB* effectively reversed the excessive biofilm phenotype conferred by *glnP* deletion (Figures 9, S18, and S19).

**Figure 9.**
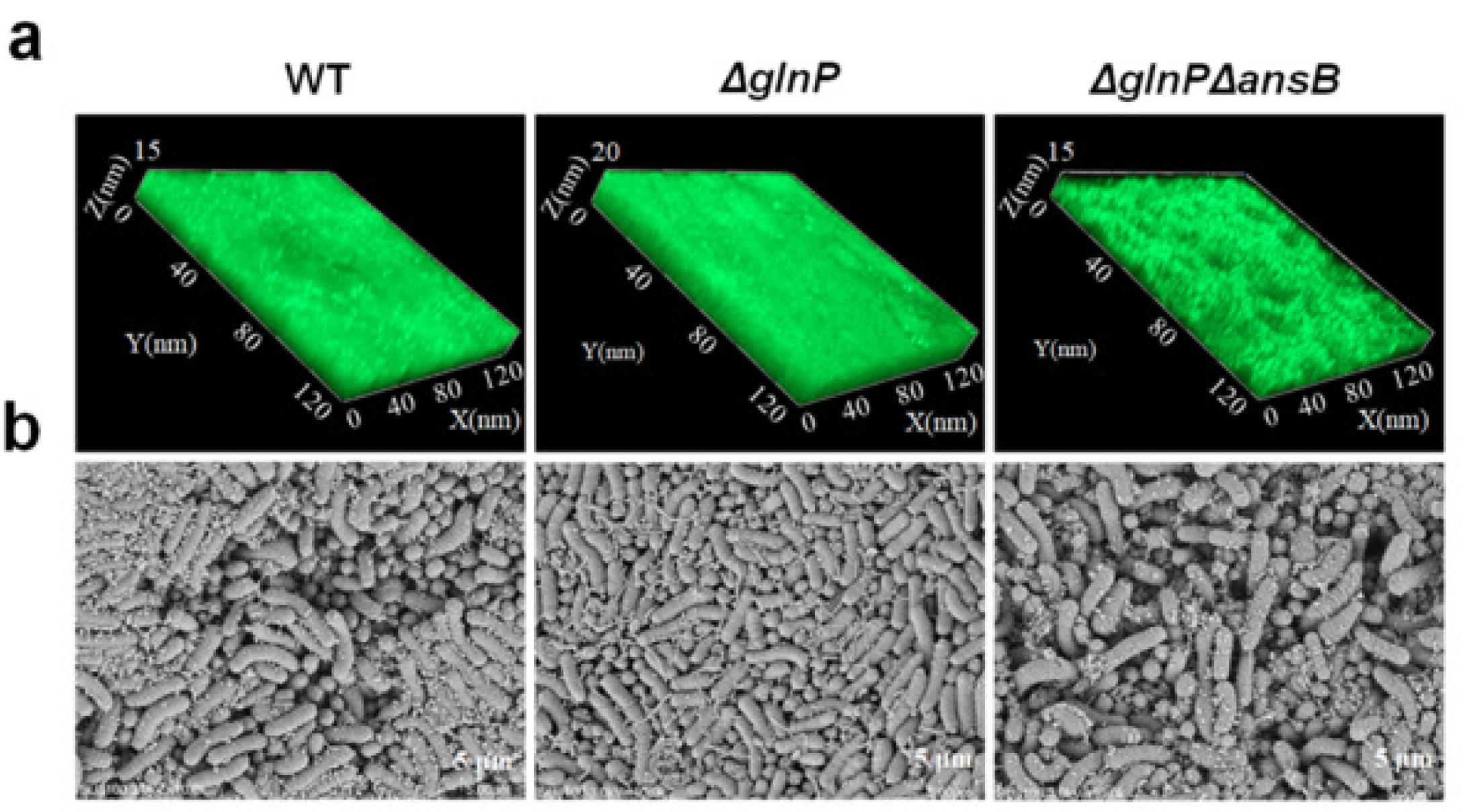
Deletion of *ansB* reverses the excessive biofilm formation phenotype of the *ΔglnP* mutant under sub-inhibitory metronidazole induction.WT, *ΔglnP,* and *ΔglnPΔansB* strains were cultured on nitrocellulose membranes containing metronidazole (1 μg/mL) for 3 days. (a) Three-dimensional CLSM reconstructions showing differences in biofilm thickness (Z-axis) among the indicated strains. (b) Representative SEM images of biofilm microarchitecture. Scale bar, 5 μm. Data are representative of three independent experiments.

### Upregulation of *ansB* induces intracellular ROS accumulation, promoting biofilm formation

To test whether *ansB* upregulation leads to increased intracellular ROS, we used the DCFH-DA fluorescent probe to monitor endogenous ROS levels in biofilm cells following sub-inhibitory antibiotic treatment. Confocal microscopy revealed intense green fluorescence in *ΔglnP* biofilm cells, whereas WT and complementation (*glnP*\*) strains displayed weak and comparable signals (Figures 10a, b). Flow cytometric quantification confirmed a rightward shift of the fluorescence peak in the *ΔglnP* strain, indicating markedly elevated ROS levels, while the *glnP*\* complementation strain reverted to WT baseline (Figure 10c). In contrast, the *ΔglnPΔansB* double mutant exhibited dramatically attenuated fluorescence, returning essentially to WT levels (Figure 11). This phenotypic rescue demonstrates that the ROS accumulation triggered by *glnP* deletion is critically dependent on *ansB* upregulation.

**Figure 10.**
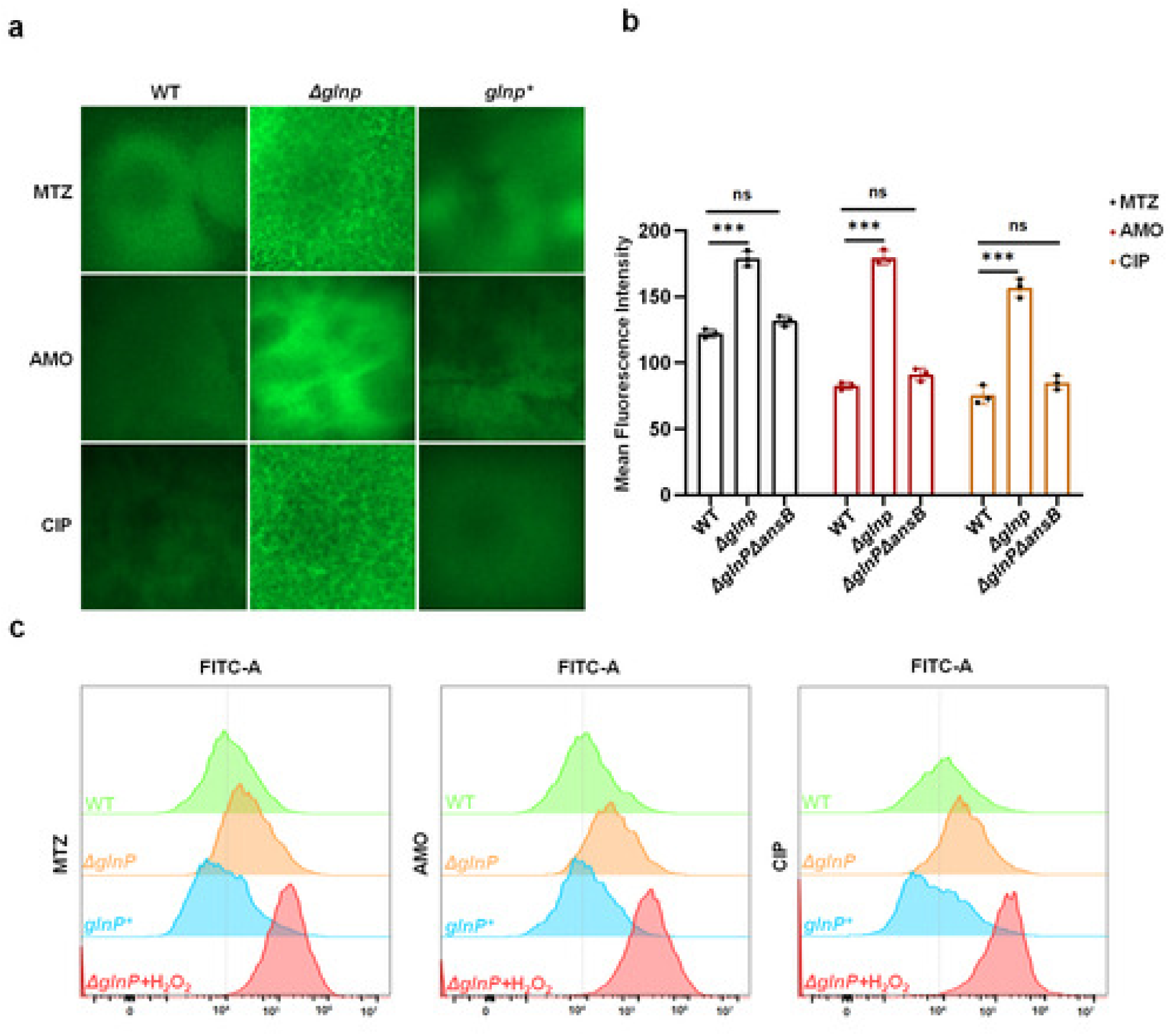
Deletion of *glnP* leads to intracellular ROS accumulation in *H. pylori* biofilms under sub-inhibitory antibiotic treatment. WT, *ΔglnP*, and *glnP\** strains were cultured with sub-inhibitory concentrations of MTZ (1 μg/mL), AMO (1/32 μg/mL), or CIP (1/16 μg/mL) for 3 days and stained with the ROS-sensitive fluorescent probe DCFH-DA. (a) Representative CLSM images showing intracellular ROS levels (green fluorescence). (b) Quantification of relative fluorescence intensity from CLSM images. (c) Flow cytometric analysis of intracellular ROS levels. Histogram peaks: green, WT; orange, *ΔglnP*; blue, *glnP*\*; red, H₂O₂ (50 μM) positive control. Data in (b) and (c) are presented as mean ± SD; ***, *P* < 0.001; ns, not significant. All experiments were performed with three independent biological replicates.

**Figure 11.**
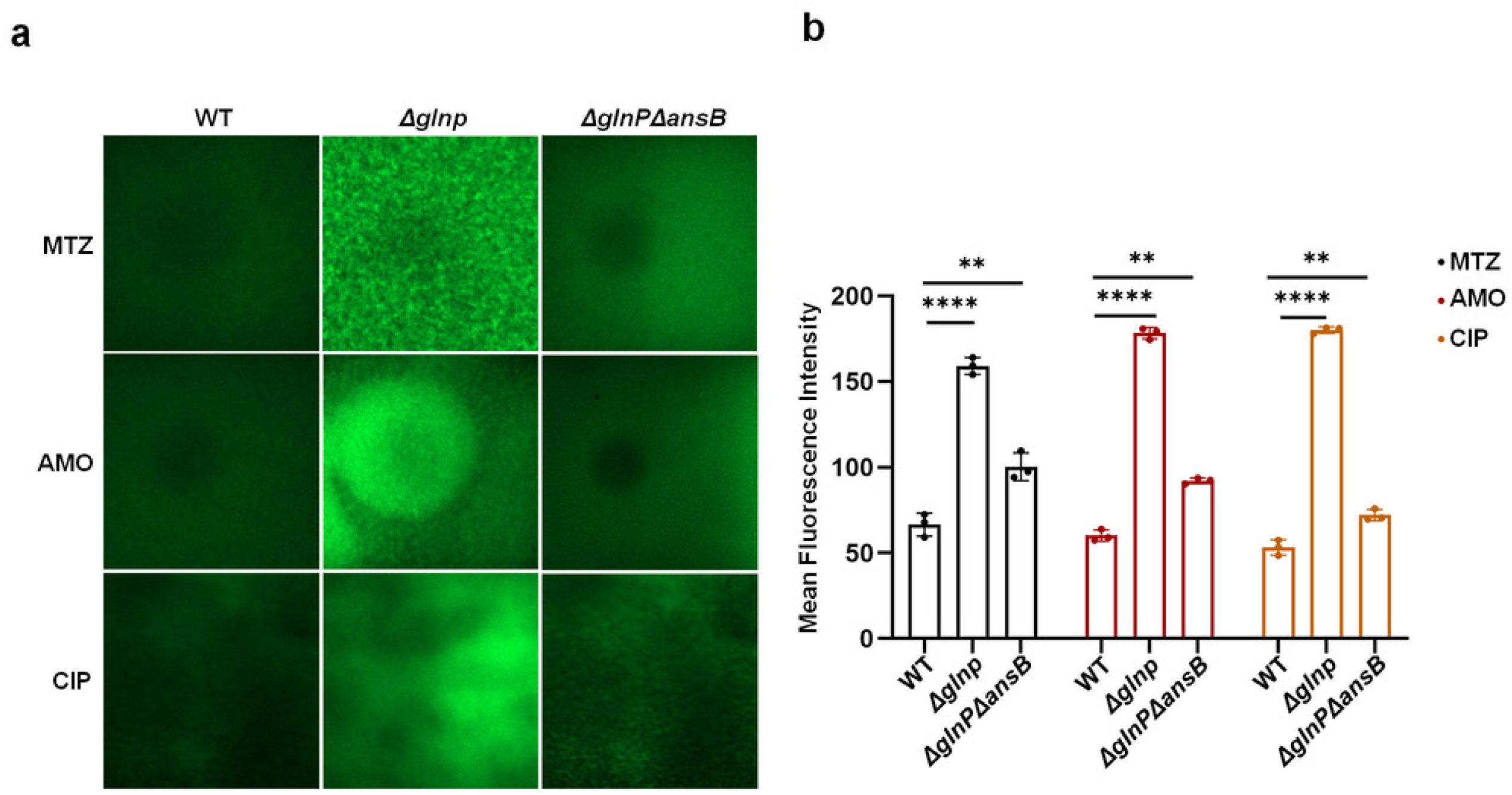
Deletion of *ansB* attenuates intracellular ROS accumulation in the *ΔglnP* mutant under sub-inhibitory antibiotic treatment. WT, *ΔglnP*, and *ΔglnPΔansB* strains were cultured with sub-inhibitory MTZ, AMO, or CIP for 3 days and stained with DCFH-DA. (a) Representative CLSM images of intracellular ROS (green fluorescence). (b) Quantification of relative fluorescence intensity from CLSM images. Data are presented as mean ± SD; **, *P* < 0.01; ****, *P* < 0.0001; ns, not significant. All experiments were performed with three independent biological replicates.

To further validate that ROS accumulation drives biofilm formation, we supplemented catalase during biofilm induction. Catalase treatment significantly inhibited biofilm formation in *ΔglnP*, *ΔglnPΔansB*, and WT strains in a concentration-dependent manner (Figure S20).

### CrdRS regulates biofilm formation by specifically responding to antibiotic signals rather than ROS

Given that two-component systems can also sense endogenous signals, we asked whether CrdRS recognizes antibiotic molecules or is activated indirectly by ROS generated under antibiotic stress. We compared biofilm formation under H₂O₂ induction between WT and the phosphorylation-defective mutants CrdR^D53A^ and CrdS^H173A^ using CLSM and SEM. All strains—including the mutants—formed normal biofilms under direct H₂O₂ exposure (Figure 12). This demonstrates that phosphorylation-site mutations in CrdRS do not impair H₂O₂-induced biofilm formation, indicating that CrdRS activation under antibiotic stress is genetically separable from ROS sensing. Thus, CrdRS specifically responds to antibiotic-induced signals rather than functioning as a downstream ROS sensor.

**Figure 12.**
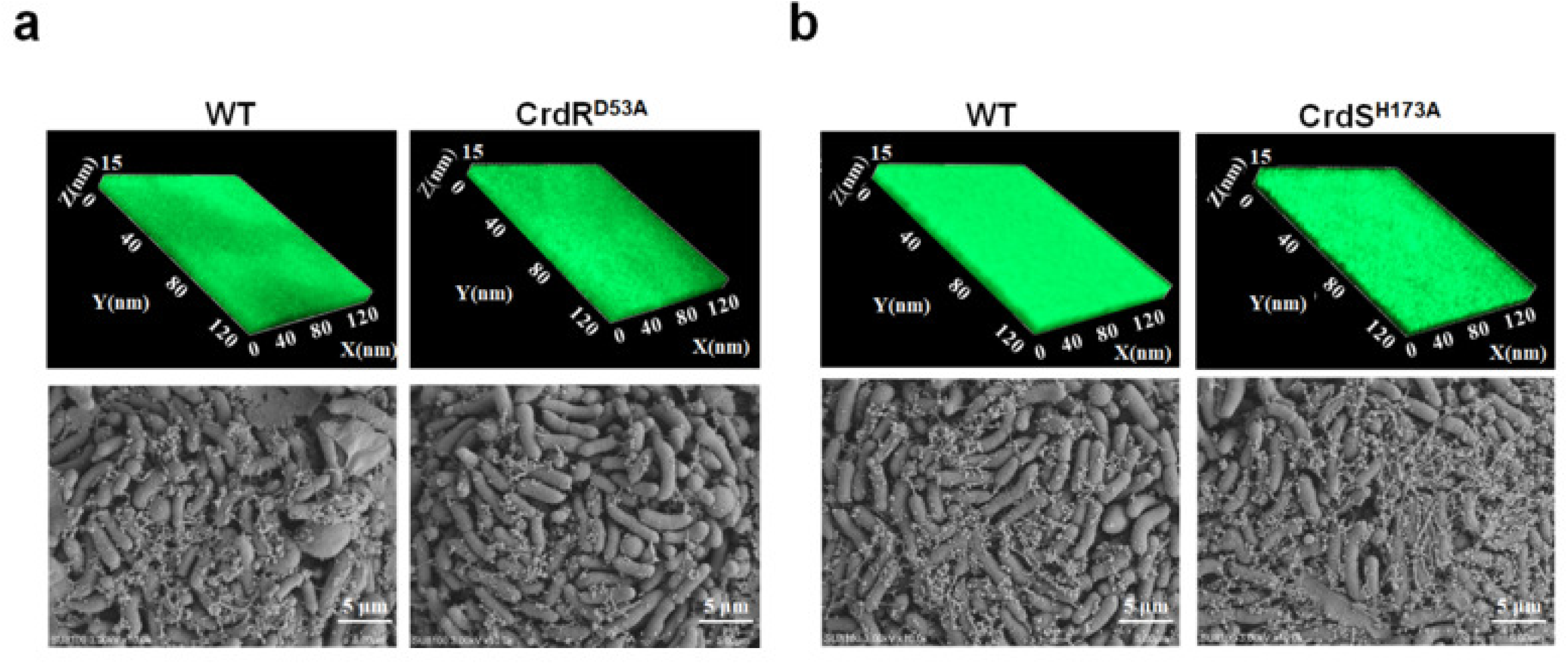
Phosphorylation-defective CrdRS mutants retain the capacity for H₂O₂-induced biofilm formation. (a, b) Biofilm morphology of WT versus CrdR^D53A^ (a) or CrdS^H173A^ (b) mutant strains following induction with 50 μM H₂O₂. Top panels: three-dimensional CLSM reconstructions showing biofilm thickness (Z-axis). Bottom panels: SEM images showing biofilm microstructure. Data are representative of three independent experiments.

### Differential expression of *glnP* and *plpA* in clinical multidrug-resistant and drug-sensitive isolates

We determined the expression levels of *glnP* and *plpA* by qRT-PCR in 8 clinical multidrug-resistant isolates and 8 clinical drug-sensitive isolates from our laboratory collection (31). For both planktonic and biofilm cells, *glnP* expression was significantly lower in multidrug-resistant strains (C-MDR) than in sensitive strains (C-DSS), whereas *plpA* showed the opposite trend, with significantly higher expression in multidrug-resistant strains (Figure 13).

**Figure 13.**
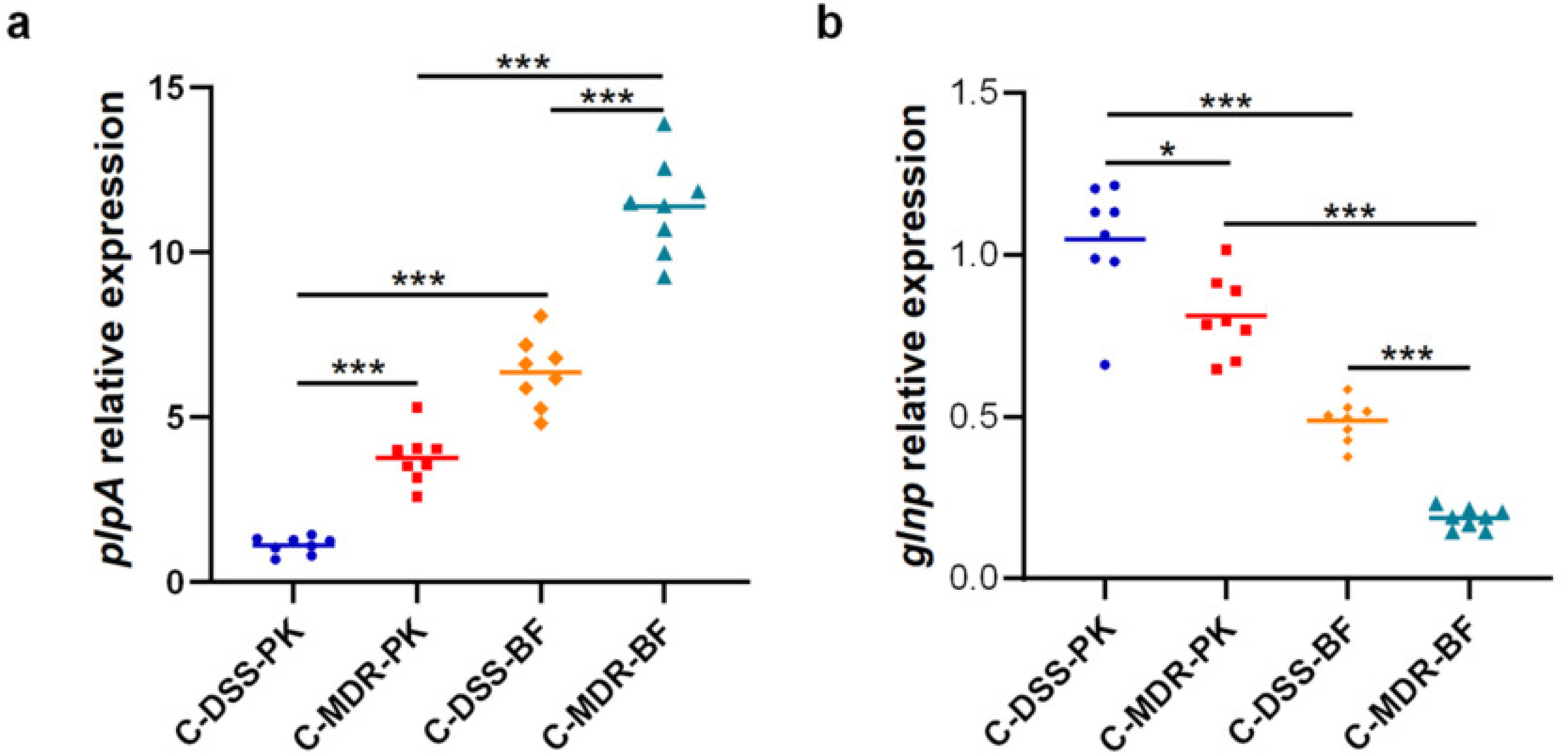
qRT-PCR analysis of *plpA* and *glnP* expression differences in clinically drug-susceptible and multidrug-resistant *H. pylori* strains. Expression levels of *plpA* and *glnP* were determined in 8 clinical drug-susceptible strains (C-DSS) and 8 clinical multidrug-resistant strains (C-MDR) under both planktonic (PK) and biofilm (BF) conditions. Results were compared to the value for *plpA* and *glnP* in planktonic WT 26695. The signal was normalized to the *16s rRNA* levels. Data are the means ± standard errors of the means from three independent experiments. Significance was determined by the paired Student’s t test. **, *P* < 0.01; ***, *P* < 0.001.

## Discussion

In the present study, we demonstrated that sub-inhibitory concentrations of clinically relevant antibiotics efficiently induce biofilm formation in *H. pylori*. Mechanistic dissection revealed that the two-component system CrdRS orchestrates this process through a dual regulatory mechanism: CrdRS positively activates the outer-membrane ABC transporter substrate-binding protein PlpA to promote exopolysaccharide secretion and matrix assembly, while concurrently repressing the inner-membrane amino acid permease GlnP to relieve transcriptional inhibition of *ansB*, triggering intracellular ROS accumulation and oxidative stress-dependent biofilm maturation. This parallel bidirectional regulation integrates distinct antibiotic stress inputs into a unified biofilm defense program.

### Sub-inhibitory concentrations of antibiotics as environmental signals for biofilm induction

Bacterial biofilm formation is not merely a passive physical aggregation, but an active biological process involving environmental sensing and transcriptional reprogramming in response to adverse external conditions (32). In both natural and host microenvironments, diverse stressors—oxidative stress (16), nutrient limitation (17), pH fluctuation (24), and antibiotic exposure (19, 20)—can be recognized by bacteria as survival signals through signaling networks such as two-component systems (33, 34), triggering stress-sensing–adaptation cascades that convert external pressure into molecular switches initiating population-protective structures (35).

From both clinical and evolutionary perspectives, sub-MIC antibiotic-induced biofilm formation is highly plausible. During eradication therapy, local drug concentrations at the infection site frequently fail to maintain stable bactericidal levels due to intragastric pharmacokinetics, mucus-layer permeability barriers, and uneven tissue distribution (36), creating exposure windows at sub-MIC concentrations. Under such conditions, antibiotics function not merely as ineffective bactericidal agents but as environmental stress signals that bacteria recognize to initiate defensive survival programs (19). A recent study by Krzyżek et al. (21) demonstrated that continuous exposure to sub-MICs of metronidazole or levofloxacin stimulated autoaggregation and EPS production in clinical H. pylori strains, resulting in increased biofilm dimensions; in contrast, clarithromycin exerted a biofilm-weakening effect, indicating that the impact of sub-MIC antibiotics on biofilm development is both antibiotic-type- and concentration-dependent. This phenomenon aligns with observations in other pathogenic bacteria: in *Staphylococcus aureus*, sub-MICs of various antibiotics modulate virulence factor expression, adherence, and biofilm formation (37, 38); in *Pseudomonas aeruginosa*, low-concentration antibiotics similarly induce biofilm development (35), suggesting that biofilm formation represents a conserved adaptive strategy for coping with sub-lethal drug pressure.

In our study, sub-inhibitory concentrations of metronidazole (1 μg/mL), amoxicillin (1/32 μg/mL), and ciprofloxacin (1/16 μg/mL) all effectively induced dense biofilm formation in *H. pylori* 26695, whereas concentrations at or above the MIC resulted in bacterial killing and failure of biofilm formation. This finding adheres to the bacterial dose-response paradigm of “low-concentration adaptation versus high-concentration death,” demonstrating that *H. pylori* possesses the capacity to sense low-concentration antibiotics and initiate biofilm-mediated defense. Notably, the three antibiotics tested target distinct cellular processes—DNA synthesis (metronidazole), cell wall synthesis (amoxicillin), and DNA gyrase (ciprofloxacin)—yet they convergently induced biofilm formation, implying a common sensing and signaling mechanism rather than drug-specific pathways. This adaptive strategy underscores the clinical importance of maintaining adequate drug concentrations throughout antibiotic therapy to avoid sub-MIC exposure windows that may inadvertently promote biofilm-associated treatment failure. Clinical evidence supports this concern: high biofilm-forming *H. pylori* isolates exhibit a significantly higher proportion of resistance to metronidazole, clarithromycin, and levofloxacin compared with low biofilm-forming isolates (39), directly linking the biofilm phenotype to clinically relevant antibiotic resistance.

### PlpA promotes biofilm matrix assembly through exopolysaccharide secretion

Bacterial biofilm formation is fundamentally dependent on the synthesis and secretion of EPS. ABC transporters participate in nutrient acquisition and solute transport, processes that sustain bacterial metabolism and indirectly support biofilm development (40). Within this system, substrate-binding proteins (SBPs) serve as critical components responsible for recognizing and capturing specific substrates—peptides, amino acids, or oligopeptides—and delivering them to cognate membrane permease complexes for transmembrane transport (41, 42).

Accumulating evidence demonstrates that functional defects in ABC transport systems significantly attenuate biofilm-forming capacity. In *Pseudomonas aeruginosa*, deletion of the substrate-binding protein DppA1 leads to an approximately 68-fold reduction in biofilm formation in static models and abolishes biofilm formation in flow cells, accompanied by reduced small-colony variant populations (40). Furthermore, overexpression of certain ABC transporter operons, such as the SppABCD system in *P. aeruginosa*, has been shown to slightly increase biofilm and exopolysaccharide synthesis (43). These findings indicate that ABC transporters influence biofilm development through diverse, yet functionally relevant, mechanisms.

In *H. pylori*, PlpA (HP1564) belongs to the ABC transporter family of substrate-binding proteins. Our study revealed that *plpA* is markedly upregulated under sub-inhibitory antibiotic induction across all three antibiotics tested, and that the *ΔplpA* mutant exhibits significantly reduced polysaccharide content in the extracellular matrix (Figure 8), leading to a structurally compromised biofilm architecture (Figure 2). The carbohydrate fraction of the *H. pylori* biofilm EPS matrix is known to be composed predominantly of mannose-related proteoglycans (proteomannans), together with fucose, galactose, N-acetylglucosamine, and 1,4-mannosyl linkages prevalent in both developing and mature biofilms (44, 45). The observation that PlpA is a periplasmic SBP suggests that it may function in recognizing and binding substrates required for EPS biosynthesis, or alternatively, that it participates in the periplasmic delivery of substrates to the inner-membrane ABC transporter complex. The near-complete loss of biofilm structural integrity in the *ΔplpA* mutant, coupled with the specific reduction in polysaccharide content, supports a model in which PlpA-mediated substrate transport constitutes a rate-limiting step for EPS matrix assembly during antibiotic-induced biofilm formation. Nevertheless, the precise substrate specificity of PlpA and the mechanism by which it contributes to exopolysaccharide secretion warrant further investigation.

### GlnP downregulation triggers *ansB*-dependent ROS accumulation to drive biofilm maturation

A particularly intriguing finding of this study is that the ABC transporter permease GlnP exerts a negative regulatory effect on *H. pylori* biofilm formation. This stands in contrast to the biofilm-promoting role of PlpA and highlights the functional diversification of ABC transporter components in biofilm regulation. The *ΔglnP* mutant formed biofilms with significantly enhanced compactness and thickness compared with WT (Figure 3), an effect that was phenocopied in the clinical isolate H57 background, confirming that GlnP deficiency promotes biofilm formation across distinct strain backgrounds.

Transcriptomic analysis provided critical mechanistic insights: the L-asparaginase gene *ansB* (*hp0723*) was markedly upregulated in the *ΔglnP* strain. AnsB catalyzes the deamination of L-asparagine to L-aspartate, which can be converted to oxaloacetate via transamination, thereby replenishing the TCA cycle intermediate pool. This metabolic flux, however, consumes α-KG as the amino group acceptor. α-KG serves as a precursor for glutamate synthesis, which is essential for GSH production—the major low-molecular-weight thiol antioxidant in bacteria—and depletion of the α-KG pool would be predicted to attenuate bacterial antioxidant defense capacity, leading to intracellular ROS accumulation(30). This prediction was experimentally validated: DCFH-DA staining and flow cytometry confirmed markedly elevated ROS in *ΔglnP* biofilm cells, whereas the *ΔglnPΔansB* double mutant showed levels comparable to WT (Figure 11). Consistently, genetic ablation of *ansB* or exogenous catalase reversed the hyper-biofilm phenotype caused by *glnP* deletion (Figures 9 and S20). Together, these rescue experiments support a model in which *glnP* downregulation derepresses *ansB*, triggering a metabolic cascade that depletes α-KG, limits glutamate for GSH synthesis, and drives ROS-dependent biofilm maturation.

Low-level ROS act as signaling molecules that promote the planktonic-to-biofilm transition in *H. pylori* (29). The ROS levels detected in the *ΔglnP* mutant likely fall within such a signaling-competent rather than cytotoxic range, as the mutant showed no growth impairment compared with WT (Figure S4b)—a hallmark of hormetic response, where low-level oxidative stress triggers adaptive protective responses (46). GlnP, as an inner-membrane permease, likely mediates uptake of amino acids whose intracellular availability influences *ansB* expression. Under steady-state conditions, GlnP-dependent transport maintains metabolic homeostasis, keeping *ansB* at basal levels; upon antibiotic stress, CrdRS-mediated repression of *glnP* disrupts this homeostasis, derepressing *ansB* and initiating ROS accumulation that drives biofilm maturation.

### CrdRS mediates antibiotic-induced biofilm formation independently of ROS sensing

The CrdRS two-component system was originally characterized as a copper-responsive regulator: CrdS senses elevated Cu²⁺ and phosphorylates CrdR, which binds AC-rich motifs (core: AACACC-ATTT-CCACAA) in the *crdA* promoter to activate copper resistance (26). Hung et al. subsequently showed that CrdRS also responds to nitrosative stress and confirmed by competitive EMSA that dual AC-rich regions are required for CrdR binding (27), establishing CrdRS as a versatile stress sensor rather than a dedicated copper-specific regulator.

This study extends the CrdRS paradigm by identifying it as a central hub for sub-inhibitory antibiotic-induced biofilm formation via bidirectional regulation of two ABC transporter genes. Two lines of evidence establish that CrdRS activation under antibiotic stress is independent of secondary ROS. First, both *plpA* and *glnP* promoters contain paired AC-rich motifs bound by phosphorylated CrdR (Figures 6 and 7). Second, phosphorylation-defective mutants CrdR^D53A^ and CrdS^H173A^ failed to form biofilms under antibiotic induction (Figure S14-S15) yet retained full biofilm-forming capacity under direct H₂O₂ exposure (Figure 12). This genetic dissociation demonstrates that CrdRS phosphorylation is required for antibiotic signal transduction but dispensable for ROS-induced biofilm formation. Whether CrdS directly binds these structurally diverse antibiotics or senses a common downstream consequence—such as membrane stress or proton motive force dissipation—warrants future biochemical and structural studies. This indirect sensing model has precedent in other bacterial stress-response systems (47) and warrants investigation using in vitro reconstitution assays.

These findings, together with recent evidence linking CrdRS to antibiotic efflux via the CrdAB-CzcBA pump (48), position CrdRS as a master regulator coordinating copper detoxification, nitrosative stress survival, antibiotic efflux, and biofilm formation through phosphorylation-dependent regulation of genes sharing AC-rich promoter elements. The dual regulatory mechanism—positive activation of PlpA for EPS production coupled with negative inhibition of GlnP for ROS generation—ensures that matrix assembly and the ROS-dependent maturation signal are produced simultaneously.

In summary, this study establishes a comprehensive model (Figure 14) in which the CrdRS two-component system serves as a central antibiotic-sensing hub that orchestrates biofilm formation in *H. pylori* through bidirectional regulation of ABC transporters, integrating extracellular matrix assembly and intracellular redox signaling into a coordinated defensive response. Importantly, the differential expression of *plpA* and *glnP* observed in clinical multidrug-resistant versus drug-susceptible isolates (Figure 13) links this CrdRS-PlpA/GlnP regulatory axis to clinically relevant antibiotic resistance. These findings identify CrdRS as a promising therapeutic target for disrupting biofilm-mediated drug tolerance and improving treatment outcomes in multidrug-resistant H. pylori infections.

**Figure 14.**
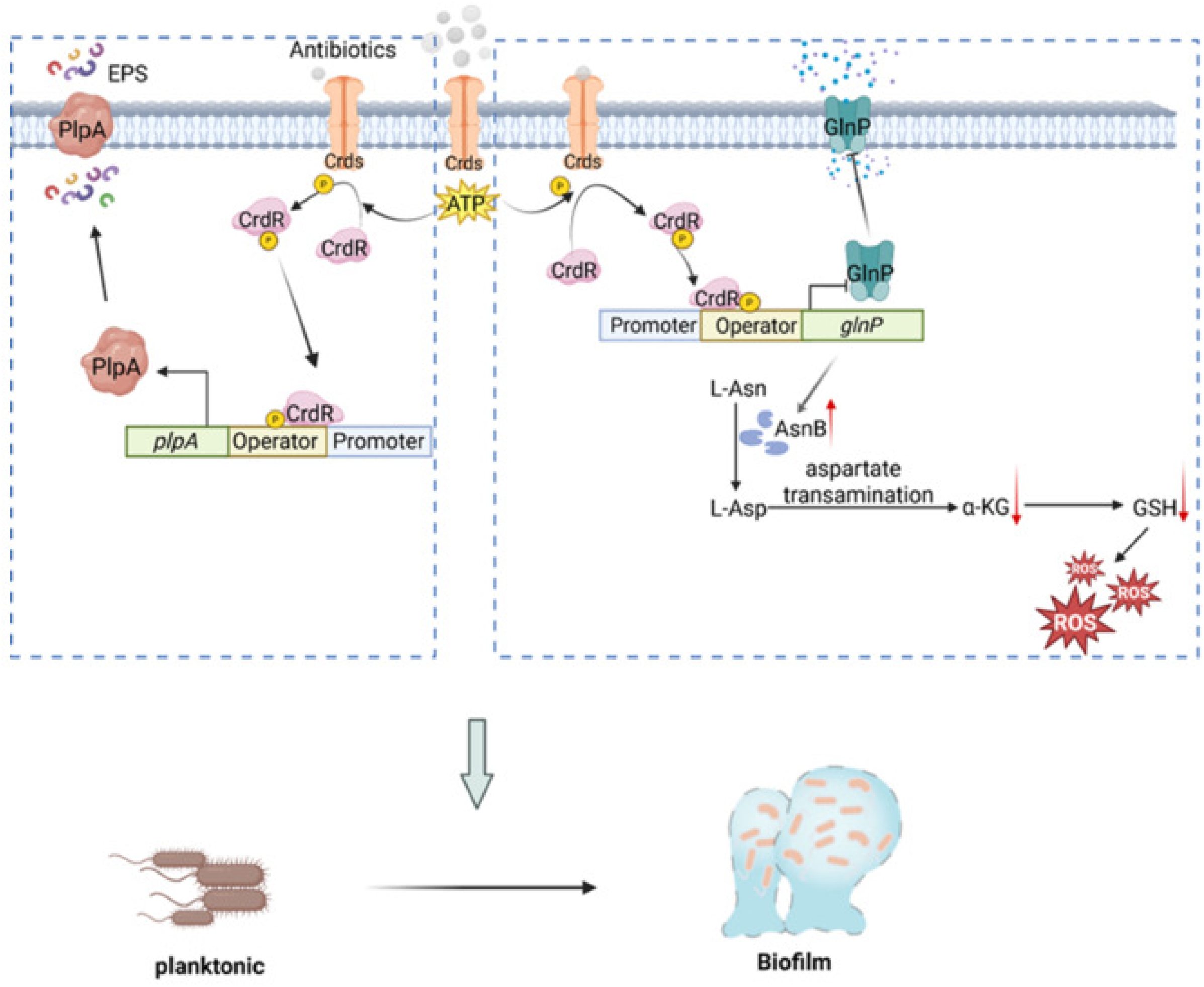
Schematic model of CrdRS-mediated regulation of sub-inhibitory antibiotic-induced biofilm formation in *H. pylori*. Upon antibiotic exposure, the histidine kinase CrdS autophosphorylates and transfers the phosphoryl group to the response regulator CrdR.Phosphorylated CrdR directly activates *plpA* transcription to promote exopolysaccharide secretion and matrix assembly (left), while concurrently repressing *glnP* transcription, leading to intracellular ROS accumulation that drives biofilm maturation (right). This dual regulatory mechanism coordinates antibiotic-induced biofilm development.

## Materials and Methods

### Bacterial Strains and Culture Conditions

*H. pylori* reference strain 26695 and clinical isolates (obtained from a gastric ulcer patient with three failed standard triple therapy courses at Qiannan Prefecture People’s Hospital, Guizhou, China) were used in this study. The study was approved by the Ethics Committee of the School of Medicine, Shandong University (Approval No. ECSBMSSDU2020-1-021). *H. pylori* was routinely cultured on Columbia agar supplemented with 5% (v/v) defibrinated sheep blood and 1% (v/v) antibiotic cocktail (polymyxin, trimethoprim, amphotericin B, vancomycin) under microaerobic conditions (5% O₂, 10% CO₂, 85% N₂) at 37°C. Liquid cultures were grown in Brucella broth containing 10% (v/v) newborn calf serum with shaking at 120 rpm in sealed gas jars. Stocks were stored in brain heart infusion with 30% (v/v) glycerol at –80°C. Escherichia coli TOP10 and BL21(DE3) were used for cloning and protein expression, respectively, and were cultured in Luria–Bertani (LB) medium at 37°C with shaking at 200 rpm. Deletion and complementation mutants were maintained with selected antibiotics: kanamycin (30 μg/mL) or chloramphenicol (100 μg/mL). All antibiotics were purchased from Sigma-Aldrich.

### Construction of *H. pylori* Gene Deletion, Complementation, and Site-Directed Mutants

Gene deletion mutants were constructed in the H. pylori 26695 background by allelic replacement with the aphA-3 kanamycin resistance cassette (49). Complementation strains were generated by inserting the target gene with its native promoter into the intergenic region between *hp0203* and *hp0204* using plasmid pBHKP252 (50), followed by natural transformation and allelic exchange. The clinical isolate H57 deletion mutants (*ΔplpA*, *ΔglnP*) were obtained using the same strategy. Phosphorylation-site mutants (CrdR^D53A^, CrdS^H173A^) were constructed by a multi-step PCR method with specific inner mutagenic primers and high-fidelity DNA polymerase. All mutants were verified by PCR and sequencing (51). Primer sequences are listed in Supplementary Table S1.

### Growth Curve Determination

Overnight *H. pylori* cultures were adjusted to an initial OD₆₀₀ of 0.08 and inoculated into Brucella broth supplemented with 10% heat-inactivated newborn calf serum. Cultures were incubated microaerobically at 37°C with shaking at 120 rpm. OD₆₀₀ was measured every 12 h for 144 h. Data are expressed as mean ± SD from three biological replicates.

### Biofilm Formation

Biofilms were cultivated using the colony biofilm assay (49). Logarithmic-phase cultures were adjusted to 1 × 10⁸ CFU/mL, and 25 μL aliquots were spotted onto nitrocellulose membranes (1 cm²) placed on solid medium. After antibiotic concentration-gradient screening, optimal sub-inhibitory induction conditions were determined as: metronidazole (1 μg/mL), amoxicillin (1/32 μg/mL), and ciprofloxacin (1/16 μg/mL). All subsequent experiments used these optimal concentrations; 50 μM H₂O₂ served as the positive control. Plates were incubated inverted under microaerobic conditions at 37°C for 2–3 days. Mature biofilms were imaged using a Zeiss stereo microscope.

### Confocal Laser Scanning Microscopy (CLSM)

Mature biofilms were gently washed with PBS to remove planktonic cells and stained with SYTO 9/propidium iodide (LIVE/DEAD Cell Viability Assay) for bacterial viability, Calcofluor White for exopolysaccharides, or SYPRO Ruby Biofilm Matrix Stain for extracellular proteins. Staining was performed in the dark at 37°C for 30 min. After washing, samples were visualized and three-dimensionally reconstructed using a Zeiss LSM 980 confocal laser scanning microscope.

### Scanning Electron Microscopy (SEM)

Biofilms were rinsed with sterile PBS and fixed with 2.5% glutaraldehyde at 4°C overnight. After washing with 0.1 M phosphate buffer (pH 7.4), samples were post-fixed in 1% osmium tetroxide for 1–2 h at room temperature, dehydrated through a graded ethanol series (30%–100%), freeze-dried, sputter-coated with gold, and examined under a Hitachi SU8100 scanning electron microscope.

### Crystal Violet Biofilm Quantification

Triplicate mature biofilms were gently washed and eluted from membranes with 1 mL PBS. One milliliter of 1% crystal violet solution was added, mixed, and incubated at room temperature for 25 min. After washing with PBS to remove excess dye, bound crystal violet was solubilized with 1 mL absolute ethanol, and OD₅₉₅ was measured using an Agilent SH1M2F Multimode Reader. Absolute ethanol served as the blank. Three biological replicates were performed.

### RNA Extraction and Quantitative Real-Time PCR (qRT-PCR)

Total RNA was extracted from *H. pylori* cells (OD₆₀₀ = 1.0) using the TRIzol method. Genomic DNA was eliminated and reverse transcription performed with the Evo M-MLV RT Mix Kit with gDNA Clean. qRT-PCR was conducted using ChamQ SYBR Color qPCR Master Mix on a CFX96 system. The 16S rRNA gene was used as the internal reference, and relative expression was calculated by the 2^−ΔΔCT^ method (52). Semi-quantitative RT-PCR products were resolved by agarose gel electrophoresis. Primer sequences are provided in Supplementary Table S1.

### Luciferase Reporter Gene Assay

The promoter regions of *glnP* and *plpA* were cloned into the pGL3-Basic vector to generate pGL3-OP*_glnP_* and pGL3-OP*_plpA_*. These reporter plasmids were co-transformed with pET32a-*crdR* into *E. coli* BL21(DE3) in various combinations. Luciferase activity was measured using a Luciferase Reporter Gene Assay Kit and a Centro XS3 LB960 luminometer. Empty pGL3-Basic and pET32a vectors served as controls. Experiments were performed in triplicate.

### Protein Expression and Purification

Wild-type *crdR* and mutant *crdR*^D53A^ (stop codon removed) were cloned into pET-32a and transformed into *E. coli* BL21(DE3). Strains were grown to OD₆₀₀≈0.6-0.8 at 37°C, induced with 0.5 mM IPTG, and incubated at 16°C with shaking at 200 rpm for 12-16 h. Recombinant proteins were purified by Ni-IDA Sepharose affinity chromatography with imidazole gradient elution. Purity was assessed by 10% SDS-PAGE and Coomassie brilliant blue staining.

### Electrophoretic Mobility Shift Assay (EMSA)

EMSA was performed as previously described (30).The *crdA* promoter (OP*_crdA_*) served as positive control (26)and the 16S rRNA gene promoter (OP*_16S_*) as negative control. Defined DNA probes encompassing conserved promoter regions were PCR-amplified (for probe details, see Figures 6e and 7e). Purified CrdR protein was subjected to in vitro phosphorylation and incubated with individual probes in binding buffer for 30 min. Binding reactions were resolved on 6% non-denaturing polyacrylamide gels in 0.25× TBE buffer (1% glycerol) at 180 V for 60 min. Phosphorylation site mutant CrdR^D53A^ was processed identically. Gels were stained with GelRed and imaged under UV transillumination.

### Reactive Oxygen Species (ROS) Detection

*H. pylori* biofilms cultured for 3 days under sub-inhibitory antibiotics were stained with 10 μM DCFH-DA in PBS at 37°C for 30 min in the dark. After washing, intracellular ROS fluorescence was detected using a Beckman Coulter CytoFLEX flow cytometer and a Zeiss LSM 980 confocal microscope (excitation 488 nm, emission 525 nm). Mean fluorescence intensity was quantified using ImageJ. All experiments were performed with at least three independent biological replicates.

### RNA Sequencing (RNA-seq) and Bioinformatic Analysis

*H. pylori* 26695 and *ΔglnP* mutant strains were cultured to logarithmic phase (OD₆₀₀ = 1.0) and inoculated into medium containing sub-inhibitory antibiotics (MTZ 1 μg/mL, AMO 1/32 μg/mL, or CIP 1/16 μg/mL) for 72 h biofilm induction. Biofilm cells were harvested, washed with PBS, snap-frozen in liquid nitrogen, and stored at −80°C. Samples were subjected to RNA-seq by Shanghai Majorbio Bio-Pharm Technology Co., Ltd. (China). Additional data mining was performed on previously generated transcriptomic datasets.

### Statistical Analysis

Data are presented as mean ± SEM from at least three independent biological replicates. Pairwise comparisons were performed using Student’s *t*-test; multiple group comparisons were analyzed by one-way ANOVA. *P* < 0.05 was considered statistically significant. GraphPad Prism version 10.6.0 was used for all statistical analyses and graphical representation.

### Data Availability

The RNA-seq data generated in this study have been deposited in the NCBI Gene Expression Omnibus (GEO) under accession numbers PRJNA1108659 and PRJNA1473680. All other data supporting the findings of this study are available within the manuscript and its supplementary materials.

## ACKNOWLEDGMENTS

This work was supported by the National Natural Science Foundation of China (grant numbers 82172313 and 82504573); the Major Scientific and Technological Innovation Project of Shandong Province (grant number 2021GXGC011305); and the Natural Science Foundation of Shandong Province (ZR2025MS1266).

## Author contributions

conceptualization—W.X.Z., L.Z., and Y.D.S.; methodology—W.X.Z., M.Z.Z., and L.Z.; investigation—W.X.Z., M.Z.Z., and L.Z., W.J.W., J.M.L., W.Y.M., X.Y.W., Z.Y.Z., H.Y., and Y.L.S.; writing (original draft)—W.X.Z.; writing (review and editing)—Y.D.S.; funding acquisition—Y.D.S. and S.J.X.; supervision—Y.D.S.

## Reference

1. Hooi JKY, Lai WY, Ng WK, Suen MMY, Underwood FE, Tanyingoh D, et al. 2017. Global prevalence of helicobacter pylori infection: Systematic review and meta-analysis. Gastroenterology 153:420–429.

2. Malfertheiner P, Megraud F, Rokkas T, Gisbert JP, Liou JM, Schulz C, et al. 2022. Management of helicobacter pylori infection: The maastricht vi/florence consensus report. Gut doi:10.1136/gutjnl-2022-327745.

3. Savoldi A, Carrara E, Graham DY, Conti M, Tacconelli E. 2018. Prevalence of antibiotic resistance in helicobacter pylori: A systematic review and meta-analysis in world health organization regions. Gastroenterology 155:1372–1382.e17.

4. Mégraud F, Graham DY, Howden CW, Trevino E, Weissfeld A, Hunt B, et al. 2023. Rates of antimicrobial resistance in helicobacter pylori isolates from clinical trial patients across the us and europe. Am J Gastroenterol 118:269–275.

5. Gisbert JP, Calvet X. 2011. Review article: The effectiveness of standard triple therapy for helicobacter pylori has not changed over the last decade, but it is not good enough. Aliment Pharmacol Ther 34:1255–68.

6. Coticchia JM, Sugawa C, Tran VR, Gurrola J, Kowalski E, Carron MA. 2006. Presence and density of helicobacter pylori biofilms in human gastric mucosa in patients with peptic ulcer disease. J Gastrointest Surg 10:883–9.

7. Yonezawa H, Osaki T, Kamiya S. 2015. Biofilm formation by helicobacter pylori and its involvement for antibiotic resistance. Biomed Res Int 2015:914791.

8. Hathroubi S, Servetas SL, Windham I, Merrell DS, Ottemann KM. 2018. Helicobacter pylori biofilm formation and its potential role in pathogenesis. Microbiol Mol Biol Rev 82.

9. Cammarota G, Branca G, Ardito F, Sanguinetti M, Ianiro G, Cianci R, et al. 2010. Biofilm demolition and antibiotic treatment to eradicate resistant helicobacter pylori: A clinical trial. Clin Gastroenterol Hepatol 8:817–820.e3.

10. Hathroubi S, Zerebinski J, Clarke A, Ottemann KM. 2020. Helicobacter pylori biofilm confers antibiotic tolerance in part via a protein-dependent mechanism. Antibiotics (Basel) 9.

11. Elshenawi Y, Hu S, Hathroubi S. 2023. Biofilm of helicobacter pylori: Life cycle, features, and treatment options. Antibiotics (Basel) 12.

12. Ilver D, Arnqvist A, Ogren J, Frick IM, Kersulyte D, Incecik ET, et al. 1998. Helicobacter pylori adhesin binding fucosylated histo-blood group antigens revealed by retagging. Science 279:373–7.

13. Mahdavi J, Sondén B, Hurtig M, Olfat FO, Forsberg L, Roche N, et al. 2002. Helicobacter pylori saba adhesin in persistent infection and chronic inflammation. Science 297:573–8.

14. Javaheri A, Kruse T, Moonens K, Mejías-Luque R, Debraekeleer A, Asche CI, et al. 2016. Helicobacter pylori adhesin hopq engages in a virulence-enhancing interaction with human ceacams. Nat Microbiol 2:16189.

15. Rader BA, Wreden C, Hicks KG, Sweeney EG, Ottemann KM, Guillemin K. 2011. Helicobacter pylori perceives the quorum-sensing molecule ai-2 as a chemorepellent via the chemoreceptor tlpb. Microbiology (Reading) 157:2445–2455.

16. Zhao Y, Cai Y, Chen Z, Li H, Xu Z, Li W, et al. 2023. Spot-mediated napa upregulation promotes oxidative stress-induced helicobacter pylori biofilm formation and confers multidrug resistance. Antimicrob Agents Chemother 65.

17. Wells DH, Gaynor EC. 2006. Helicobacter pylori initiates the stringent response upon nutrient and ph downshift. J Bacteriol 188:3726–9.

18. Hou C, Yin F, Wang S, Zhao A, Li Y, Liu Y. 2022. Helicobacter pylori biofilm-related drug resistance and new developments in its anti-biofilm agents. Infect Drug Resist 15:1561–1571.

19. Davies J, Spiegelman GB, Yim G. 2006. The world of subinhibitory antibiotic concentrations. Curr Opin Microbiol 9:445–53.

20. Andersson DI, Hughes D. 2014. Microbiological effects of sublethal levels of antibiotics. Nat Rev Microbiol 12:465–78.

21. Krzyżek P, Migdał P, Tusiewicz K, Zawadzki M, Szpot P. 2024. Subinhibitory concentrations of antibiotics affect development and parameters of helicobacter pylori biofilm. Front Pharmacol 15:1477317.

22. Hoffman LR, D’Argenio DA, MacCoss MJ, Zhang Z, Jones RA, Miller SI. 2005. Aminoglycoside antibiotics induce bacterial biofilm formation. Nature 436:1171–5.

23. Kaplan JB. 2011. Antibiotic-induced biofilm formation. Int J Artif Organs 34:737–51.

24. Pflock M, Finsterer N, Joseph B, Mollenkopf H, Meyer TF, Beier D. 2006. Characterization of the arsrs regulon of helicobacter pylori, involved in acid adaptation. J Bacteriol 188:3449–62.

25. Spohn G, Scarlato V. 1999. Motility of helicobacter pylori is coordinately regulated by the transcriptional activator flgr, an ntrc homolog. J Bacteriol 181:593–9.

26. Waidner B, Melchers K, Stähler FN, Kist M, Bereswill S. 2005. The helicobacter pylori crdrs two-component regulation system (hp1364/hp1365) is required for copper-mediated induction of the copper resistance determinant crda. J Bacteriol 187:4683–8.

27. Hung CL, Cheng HH, Hsieh WC, Tsai ZT, Tsai HK, Chu CH, et al. 2015. The crdrs two-component system in helicobacter pylori responds to nitrosative stress. Mol Microbiol 97:1128–41.

28. Servetas SL, Carpenter BM, Haley KP, Gilbreath JJ, Gaddy JA, Merrell DS. 2016. Characterization of key helicobacter pylori regulators identifies a role for arsrs in biofilm formation. J Bacteriol 198:2536–48.

29. Zheng Y, Li S, Xue J, Zhang L, Wang L, Zhao Y, et al. 2025. Ros-induced allosteric modulation of nikr promotes helicobacter pylori biofilm formation by attenuating flgr-dependent inhibition of the molybdate transport system. Virulence 16:2589562.

30. Leduc D, Gallaud J, Stingl K, de Reuse H. 2010. Coupled amino acid deamidase-transport systems essential for helicobacter pylori colonization. Infect Immun 78:2782–92.

31. Ge X, Cai Y, Chen Z, Gao S, Geng X, Li Y, et al. 2018. Bifunctional enzyme spot is involved in biofilm formation of helicobacter pylori with multidrug resistance by upregulating efflux pump hp1174 (glup). Antimicrob Agents Chemother 62.

32. Hathroubi S, Hu S, Ottemann KM. 2020. Genetic requirements and transcriptomics of helicobacter pylori biofilm formation on abiotic and biotic surfaces. NPJ Biofilms Microbiomes 6:56.

33. Mitrophanov AY, Groisman EA. 2008. Signal integration in bacterial two-component regulatory systems. Genes Dev 22:2601–11.

34. Alvarez AF, Georgellis D. 2023. Environmental adaptation and diversification of bacterial two-component systems. Curr Opin Microbiol 76:102399.

35. Luo Y, Zhao K, Baker AE, Kuchma SL, Coggan KA, Wolfgang MC, et al. 2015. A hierarchical cascade of second messengers regulates pseudomonas aeruginosa surface behaviors. mBio 6.

36. Nakamura M, Spiller RC, Barrett DA, Wibawa JI, Kumagai N, Tsuchimoto K, et al. 2003. Gastric juice, gastric tissue and blood antibiotic concentrations following omeprazole, amoxicillin and clarithromycin triple therapy. Helicobacter 8:294–9.

37. Kaplan JB, Izano EA, Gopal P, Karwacki MT, Kim S, Bose JL, et al. 2012. Low levels of β-lactam antibiotics induce extracellular DNA release and biofilm formation in staphylococcus aureus. mBio 3:e00198–12.

38. Haddadin RN, Saleh S, Al-Adham IS, Buultjens TE, Collier PJ. 2010. The effect of subminimal inhibitory concentrations of antibiotics on virulence factors expressed by staphylococcus aureus biofilms. J Appl Microbiol 108:1281–91.

39. Tao W, Ma W, Zhao G, Zhang Z, Xu C, Wang P, et al. 2025. Characterization of antibiotic resistance and biofilm formation in clinical helicobacter pylori isolates from ningxia, china. Sci Rep 15:45439.

40. Lee Y, Song S, Sheng L, Zhu L, Kim JS, Wood TK. 2018. Substrate binding protein dppa1 of abc transporter dppbcdf increases biofilm formation in pseudomonas aeruginosa by inhibiting pf5 prophage lysis. Front Microbiol 9:30.

41. Berntsson RP, Smits SH, Schmitt L, Slotboom DJ, Poolman B. 2010. A structural classification of substrate-binding proteins. FEBS Lett 584:2606–17.

42. Tam R, Saier MH, Jr. 1993. Structural, functional, and evolutionary relationships among extracellular solute-binding receptors of bacteria. Microbiol Rev 57:320–46.

43. Pletzer D, Braun Y, Weingart H. 2016. Swarming motility is modulated by expression of the putative xenosiderophore transporter sppr-sppabcd in pseudomonas aeruginosa pa14. Antonie Van Leeuwenhoek 109:737–53.

44. Yang FL, Hassanbhai AM, Chen HY, Huang ZY, Lin TL, Wu SH, et al. 2011. Proteomannans in biofilm of helicobacter pylori atcc 43504. Helicobacter 16:89–98.

45. Stark RM, Gerwig GJ, Pitman RS, Potts LF, Williams NA, Greenman J, et al. 1999. Biofilm formation by helicobacter pylori. Lett Appl Microbiol 28:121–6.

46. Cáp M, Váchová L, Palková Z. 2012. Reactive oxygen species in the signaling and adaptation of multicellular microbial communities. Oxid Med Cell Longev 2012:976753.

47. Bader MW, Sanowar S, Daley ME, Schneider AR, Cho U, Xu W, et al. 2005. Recognition of antimicrobial peptides by a bacterial sensor kinase. Cell 122:461–72.

48. Gao F, Xiang W, Zhang X, Huang X, She F, Wen Y. 2025. Copper enhances tetracycline resistance via the efflux transporter crdab-czcba in helicobacter pylori. Front Med (Lausanne) 12:1552537.

49. Zhao Y, Chen Z, Cai Y, Xue J, Zhang L, Wang L, et al. 2024. Aloe-emodin destroys the biofilm of helicobacter pylori by targeting the outer membrane protein 6. Microbiol Res 278:127539.

50. Bi H, Zhu L, Jia J, Cronan JE. 2016. A biotin biosynthesis gene restricted to helicobacter. Sci Rep 6:21162.

51. Wang L, Wang S, Zhang W, Zheng Y, Liu J, Ma W, et al. 2025. Ros-induced allosteric regulation of nikr coordinates hp0910-mediated omp2 methylation to modulate h. Pylori biofilm dynamics and therapeutic targeting. Microbiol Res 301:128319.

52. Livak KJ, Schmittgen TD. 2001. Analysis of relative gene expression data using real-time quantitative pcr and the 2(-delta delta c(t)) method. Methods 25:402–8.

